# Mitochondrial ribosomal proteins developed unconventional mitochondrial targeting signals due to structural constraints

**DOI:** 10.1101/2021.04.26.441393

**Authors:** Yury S. Bykov, Tamara Flohr, Felix Boos, Johannes M. Herrmann, Maya Schuldiner

**Affiliations:** Department of Molecular Genetics, Weizmann Institute of Science, Rehovot, Israel; Division of Cell Biology, University of Kaiserslautern, Germany; Department of Genetics, Stanford University, Stanford, CA

**Keywords:** mitochondria, targeting, translocation, mitochondrial ribosome, mitochondrial targeting signal, ribosome structure

## Abstract

Mitochondrial ribosomes are complex molecular machines indispensable for respiration. Their assembly involves the import of several dozens of mitochondrial ribosomal proteins (MRPs), encoded in the nuclear genome, into the mitochondrial matrix. Available proteomic and structural data as well as computational predictions indicate that up to 25% of MRPs do not have a conventional N-terminal mitochondrial targeting signal (MTS). We characterized a set of 15 yeast MRPs *in vivo* and showed that 30% of them use internal mitochondrial targeting signals. We isolated a novel internal targeting signal from the conserved MRP Mrp17 (bS6). The Mrp17 targeting signal shares some properties as well as import components with conventional MTS-containing proteins but is not reliably predicted indicating that mitochondrial protein targeting is more versatile than expected. We hypothesize that internal targeting signals arose in MRPs when the N-terminus extension was constrained by ribosome assembly interfaces.

## Introduction

Mitochondria are descendants of ancient bacteria that formed eukaryotic cells together with their archaeal host (Martijn et al., 2018; Sagan, 1967; Zaremba-Niedzwiedzka et al., 2017). Since then, mitochondria have lost their autonomy and their reproduction depends entirely on the nuclear genome, which encodes the majority of mitochondrial proteins. However all mitochondria capable of respiration have retained small vestigial genomes of their own and fully functional gene expression machineries of bacterial origin (Roger et al., 2017). Mitochondrial ribosomes (mitoribosomes) are the most complex components of the mitochondrial gene expression system and consist of several RNA molecules and 60 to 80 different proteins (Greber and Ban, 2016; Ott et al., 2016). Mitoribosome dysfunction has adverse consequences leading to a broad spectrum of diseases (Boczonadi and Horvath, 2014).

While it took many years to solve the first ribosome structures (Ban et al., 2000; Carter et al., 2000; Schluenzen et al., 2000), rapid progress in cryo-electron microscopy is now rapidly revealing the structural details of mitoribosomes of many different organismic groups (Amunts et al., 2015; Desai et al., 2017; Itoh et al., 2020; Kummer et al., 2018; Ramrath et al., 2018; Tobiasson and Amunts, 2020; Waltz et al., 2020). The availability of so many structures highlighted an interesting feature of mitoribosomes - their incredible evolutionary diversity (Kummer and Ban, 2021; Waltz and Giegé, 2019). The composition of mitochondrial ribosomes in different eukaryotic lineages underwent dramatic changes caused by multiple losses of RNA segments and mitoribosomal proteins (MRPs) as well as acquisition of new, lineage-specific RNA segments and MRPs (Desmond et al., 2011; Petrov et al., 2019; Sluis et al., 2015; Smits et al., 2007). As a result, mitoribosomes contain a core set of MRPs homologous to the bacterial ribosomal proteins (BRPs) and a variable set of MRPs that can be common for all mitochondrial ribosomes or specific only to certain eukaryotic lineages. In addition, during their evolution many MRPs acquired significant expansions of their C- and N-termini while retaining structurally conserved domains of their BRP ancestors (Melnikov et al., 2018; Sluis et al., 2015; Vishwanath et al., 2004).

Mitochondrial genomes in many eukaryotic cells (with the exception of animals) still contain genes for a number of ribosomal proteins, indicating that their successful transfer to the nuclear genome might be less easily feasible than that of many other matrix proteins (Bertgen et al., 2020). However, most, in animals even all, MRPs are nuclear encoded. Thus, similarly to the majority of mitochondrial proteins (numbering from around 800 in yeast to around 1500 in mammals), they must be imported from the cytosol (Morgenstern et al., 2017; Pagliarini et al., 2008; Vögtle et al., 2017). The import of mitochondrial proteins can be conceptually subdivided in two steps: (1) targeting of the newly synthesized mitochondrial protein precursors to the mitochondrial membrane. This can occur either post-translationally or co-translationally involving ribosome-nascent chain complexes. (2) Translocation of the unfolded precursors through the mitochondrial membrane(s) to deliver them to their final destination within mitochondria (Bykov et al., 2020). Effective targeting and translocation are mediated by specialized protein complexes that recognize targeting and translocation signals within precursor protein sequences.

Most matrix and inner membrane proteins are synthesized with N-terminal matrix targeting sequences (MTSs), also called presequences, which are both necessary and sufficient for mitochondrial targeting. MTSs have a characteristic structure that can be predicted by prediction algorithms (Armenteros et al., 2019; Claros and Vincens, 1996; Emanuelsson et al., 2000; Fukasawa et al., 2015). MTSs are typically between 10 and 60 residues in length and can form an amphipathic *α*-helix with one positively charged surface and one hydrophobic surface. In most cases MTSs are proteolytically removed during protein import, giving rise to mature forms of mitochondrial matrix or inner-membrane proteins (Bedwell et al., 1989; Vögtle et al., 2009; von Heijne, 1986).

In contrast to basically all other proteins of the mitochondrial matrix, many MRPs lack N-terminal MTSs (Woellhaf et al., 2014). In some cases, MRPs use N-terminal regions that mimic the properties of MTSs but are not cleaved (un-cleaved MTSs). Such un-cleaved MTSs are also found in some matrix proteins that are not associated with the ribosome, such as Hsp10 (Poveda-Huertes et al., 2020). Surprisingly, a number of MRPs do not contain any regions that show MTS-like features and it is unknown how mitochondria recognize and import these proteins. For now, there are only two well characterized examples of MRPs with unconventional MTSs - Mrpl32 (bL32, by new nomenclature (Ban et al., 2014)) and Mrp10 (mS37) whose import path deviates from the canonical matrix-targeting route (Bonn et al., 2011; Longen et al., 2014; Nolden et al., 2005).

In this study, we studied the mechanisms by which MRPs are imported and assembled into the mitoribosome. We systematically examined N-termini of unconventional MRPs and analyzed them *in silico* and experimentally. We further focused on the biogenesis of Mrp17 (bS6) as a representative of the unconventional group of MTS-less MRPs. We discovered a novel mitochondrial matrix targeting region that is displayed in the internal sequence of the protein. This stretch shares properties with mitochondrial targeting sequences such as positive charges for receptor binding and membrane potential-dependent translocation, but differs in its structural features and position in the protein. The efficient import of Mrp17 shows that the mitochondrial import machinery is much more versatile in its substrate spectrum than expected. More generally, our work shows how structural restrictions forced the generation of unconventional targeting motifs.

## Results

### Mapping unconventional MRP targeting signals

To systematically investigate MRP targeting signals in detail, we compiled all existing data on the maturation of their N-termini in yeast (Table S1). We used direct N-terminal sequencing data (Boguta et al., 1992; Dang and Ellis, 1990; Davis et al., 1992; Graack et al., 1988; Graack et al., 1991; Grohmann et al., 1989; Grohmann et al., 1991; Kitakawa et al., 1990; Kitakawa et al., 1997; Matsushita and Isono, 1993; Matsushita et al., 1989), N-terminal proteomics (Vögtle et al., 2009) and predictions performed by UniProt annotators, as well as by ourselves using MitoFates for cleavage site prediction (Fukasawa et al., 2015). Importantly, we also used available structural information (Desai et al., 2017). In particular, mitoribosome structures were helpful to identify proteins that do not have a cleavable MTS - such proteins had their N-termini contained within the structure and hence could not have been cleaved after import into the mitochondrial matrix. We reanalyzed ribosome profiling data on translation initiation in yeast (see Methods for details) to ascertain that none of these proteins has mis-annotated translation start sites that might produce an N-terminal extension accounting for a missing cleavable MTS (Fig. S1). In the yeast mitochondrial ribosome structure (PDB:5MRC), the structures of six proteins started with amino acid number 1 (Met), structures of 12 started with amino acid number 2, five - with amino acids 3 to 9, and the rest, 50, with amino acid number 10 and more. The number of the first amino acid present in the structure was moderately conserved among the determined mitoribosome structures (Fig. S2) and was not restricted to any particular group of MRPs classified by origin (bacterial, mitochondria-specific, or yeast-specific) or position in the structure (Fig. S3).

Interestingly, a simple distinction by the first amino acid appearing in the structure separates MRPs into two classes. In the first group are those MRPs that are derived from cleaved precursors (which consistently have high MTS prediction scores). In addition, this group may contain proteins with an uncleavable N-terminus of a flexible nature which would then be unresolved in the available structures. Some of the latter may have poor mitochondrial targeting scores in prediction algorithms. In the second group are those whose structure starts with amino acid number less than 10. Most of these proteins score very poorly with different software predicting N-terminal MTS (Fig. 1A, Fig. S2). Many MRPs of this group lack conventional, N-terminal import signals, and their targeting signals are not predictable by available software. Thus, the available structures of mitochondrial ribosomes confirm the previous conclusion that many MRPs are made without N-terminal MTSs (Woellhaf et al., 2014).

**Fig. 1.**
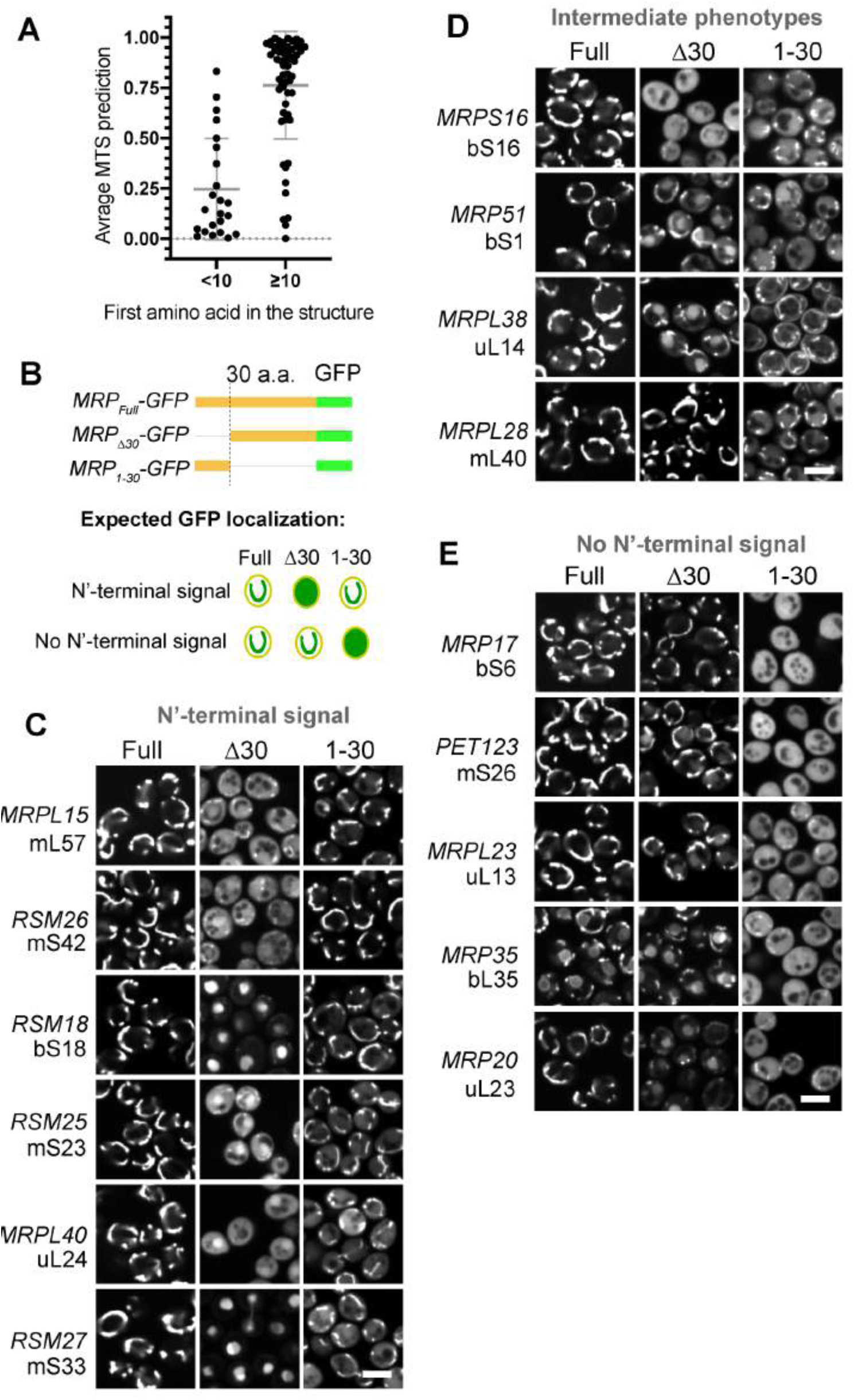
Mitochondrial Ribosomal Proteins (MRPs) have various types of targeting signals. (A) Yeast MRPs having uncleaved N-termini that can be tracked in the mitoribosome structure (PDB:5MRC) score much lower with MTS prediction algorithms (average of TargetP2 and MitoFates) compared to other MRPs that have their N-termini cleaved off or are not present in the structure (so might be flexible and outside the mitoribosome body). (B) Schematic of MRP truncations used to characterize targeting properties of MRP N-termini: MRPFull as control, MRPL\30 to check if the N-terminus is necessary, MRP1-30 to check if it is sufficient (top) and the schematics of expected GFP localization in case the N-terminus is MTS-like (necessary and sufficient) or not (bottom); (C)-(E) Micrographs collected in the GFP channel for each truncation (columns) of each studied MRP (rows) grouped by the N-terminus targeting properties based on theoretical expectation summarized in (B) with the MRPs possessing MTS-like N-termini in panel (C), MRPs with intermediate phenotype in panel (D) and MRPs without N-terminal signal in panel (D), for each MRP a yeast gene name and new nomenclature protein name is shown on the left. Scale bar for all micrographs is 5 µm.

Next, we experimentally analyzed the targeting information in the sequences of different MRPs by GFP fusion proteins. To this end, we selected 15 MRPs with different properties (Fig. S3, Table S2). Then we tested whether the N-terminal 30 residues of these proteins were necessary and/or sufficient for mitochondrial targeting. The length of 30 residues was chosen as it corresponds to the most common size of a cleavable yeast MTS (Vögtle et al., 2009). To test this, we expressed each MRP in diploid yeast fused to GFP. To assay if the N-terminus is necessary we expressed a truncated version with the first 30 amino acids deleted (MRP_Δ30_-GFP). To test if the N-terminus is sufficient we expressed a version with only the first 30 amino acids (MRP_1-30_-GFP). As a control we used the full-length version (MRP_Full_-GFP) (Fig. 1B). The distribution of GFP signals was imaged in cells in which mitochondria were stained with MitoTracker Orange (Fig. 1C-E, Fig S4, Table S2). Six proteins (Mrpl15, Rsm26, Rsm18, Rsm25, Mrpl40, and Rsm27) contained targeting information within their N-termini (Fig. 1C); of them, only Mrpl15 (mL57) had high MTS prediction scores consistent with highly confident annotation of a cleavable 29-amino acid long MTS (Table S1). Other proteins whose N-termini were able to target GFP to mitochondria had low MTS prediction scores (Table S2) indicating that their N-terminal signals have distinct properties, not similar to conventional MTSs. For four proteins (Mrps16, Mrp51, Mrpl38, and Mrpl28) neither the N-terminal 30 residues nor the internal segment on its own were sufficient for targeting, indicating that the necessary targeting information is contained in an N-terminal segment longer than 30 amino acids or distributed over the whole length of these proteins (Fig. 1D). Finally, five proteins (Mrp17, Pet123, Mrpl23, Mrp35, and Mrp20) were targeted to mitochondria independently of their N-terminal regions indicating that the targeting signals in these proteins are internal (Fig. 1E).

Interestingly, many of the N-terminally truncated MRP versions accumulated outside of the mitochondria in the cytosol or, in many cases, in the nucleus (Fig. 1C-E, Fig. S4B). These observations agree with the recent discovery that mistargeted mitochondrial proteins can accumulate in the nucleus and get degraded in perinuclear puncta (Shakya et al., 2021). Despite the mislocalization of several of these forms, none of them resulted in obvious growth defects (Fig. S5).

To summarize, we selected a subset of MRPs with diverse structural and sequence features and characterized the mitochondrial targeting capacity of their N termini. We observed that many of these MRPs contain multiple unconventional targeting signals, often outside of the 30 N-terminal residues, thus apparently scattered over their sequence. One particularly intriguing MRP was Mrp17 (bS6), a protein of the small subunit of the yeast mitoribosome. Mrp17 lacks any identifiable targeting signal while being one of the most structurally conserved proteins across all mitoribosomal structures studied to date. Hence, we chose Mrp17 for further investigation.

### Defining Mrp17 targeting and translocation signals

To investigate the unconventional mitochondrial targeting signals of Mrp17 in more detail we created a systematic set of Mrp17 truncations fused to GFP and expressed them in diploid yeast (Fig. S6). We observed that the internal fragment of Mrp17 between amino acids 20 and 100 was the minimal fragment able to target GFP to mitochondria similarly to full-length Mrp17 (131 amino acids) without producing cytosolic background signal (Fig. 2A). This indicates that similarly to the N-terminus, the C-terminus is dispensable for targeting. Splitting this fragment in two halves showed that the N-terminal part (Mrp17_21-60_) was still able to target GFP to mitochondria although with significant cytosolic background while the C-terminal part (Mrp17_61-100_) was cytosolic (Fig. 2A). We conclude that, *in vivo,* Mrp17 region 21-60 is necessary for mitochondrial targeting but is not sufficient for efficient targeting, which is promoted by additional signals distributed over the whole length of the protein (Fig. S7A,B).

**Fig. 2.**
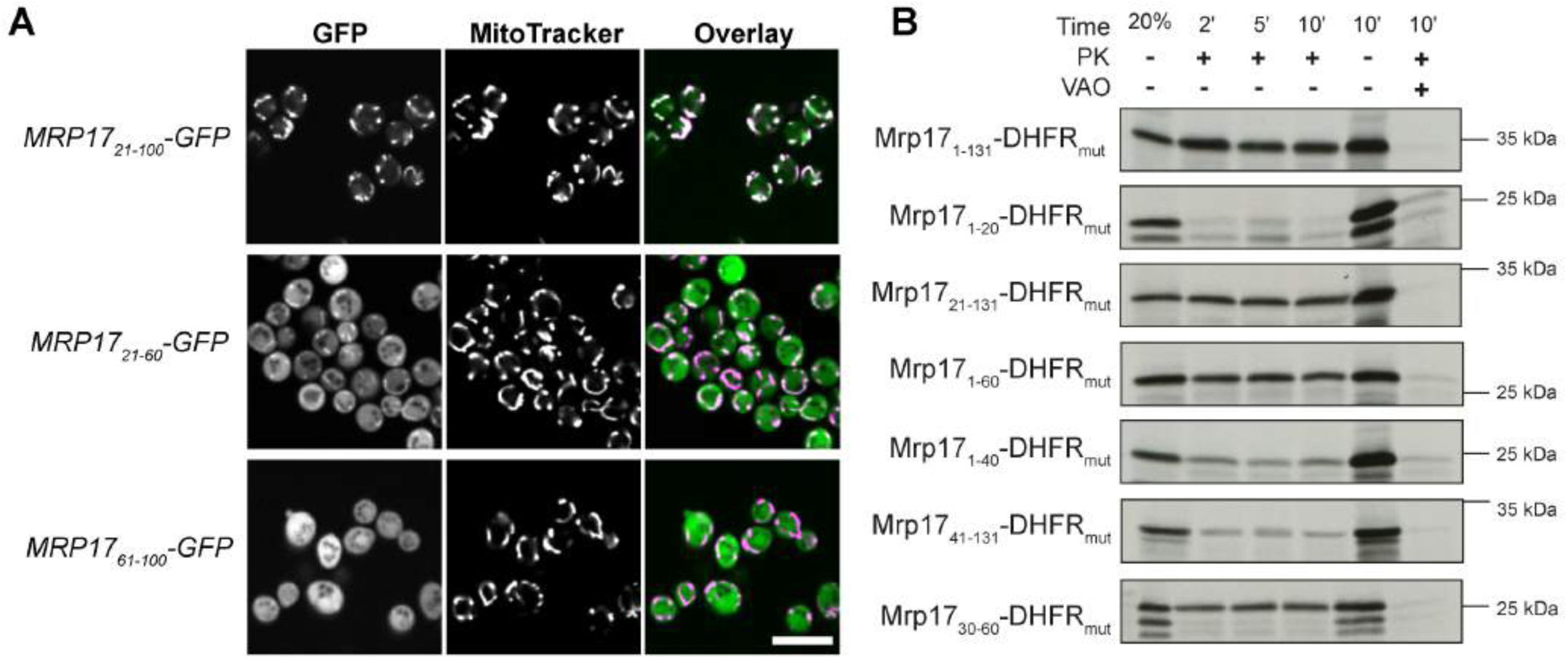
The non-canonical targeting and translocation signal of Mrp17 is located between amino acids 30 and 60. (A) In vivo characterization of mitochondrial targeting capacity of different Mrp17truncationsfused to GFP visualized by fluorescent microscopy with MitoTracker Orange straining. Scale bar for all micrographs is 10 µm. (B) Characterization of Mrp17 translocation signal using an in vitro import assay: shown are autoradiographs of full-length Mrp17 or its truncations fused to DHFRmut, translated in vitro with radiolabeled amino-acids, incubated with isolated yeast mitochondria for 2, 5, or 10 min, treated with proteinase K (PK) to remove nonimported proteins and visualized by SDS-PAGE/autoradiograph.y As a negative control, mitochondria were treated with valinomycin, antimycin and oligomycin (VAO) that eliminate membrane potential. For comparison, 20% of the protein used per import reaction was loaded on the first lane.

The microscopic analysis does not allow us to discriminate between targeting to the mitochondrial surface from complete translocation into the matrix. To elucidate the import efficiency of different Mrp17 regions we used *in vitro* translocation assays into isolated yeast mitochondria. Since Mrp17 is very small and many fragments lacked methionine residues that are necessary for radiolabeling, we fused Mrp17 to an unfolded mutant of the mouse dihydrofolate reductase - DHFR_mut_ (Vestweber and Schatz, 1988). The full-length Mrp17-DHFR_mut_ fusion was effectively imported into isolated yeast mitochondria at the same rate as untagged Mrp17 but gave much stronger signals in autoradiography (Fig. 2B, Fig. S7C). In agreement with the targeting experiments performed *in vivo*, the short N-terminal region of Mrp17 was neither necessary nor sufficient for efficient translocation (Mrp17_21-131_, Mrp17_1-20_ in Fig. 2B). The first 60 amino acids of Mrp17 were sufficient for translocation narrowing down the import signal to the N-terminal half of the protein (Mrp17_1-60_ in Fig. 2B). Leaving only the first 40 amino acids or removing them from the N-terminus reduced the translocation speed indicating that regions 20-40 and 40-60 are equally important parts of the signal (Mrp17_1-40_ and Mrp17_41-131_ in Fig. 2B). Finally, we narrowed down the Mrp17 region containing the translocation signal to amino acids 30-60 (Fig. 2B, bottom; Fig. S7D). However, similarly to the results of *in vivo* experiments, even short fragments of Mrp17 outside this region retained some translocation capacity (Fig. 2B, Fig. S7C).

To summarize, we determined that the main mitochondrial targeting and translocation signal of Mrp17 is positioned between amino acids 30 to 60. However, there exist additional signals that improve mitochondrial targeting efficiency or stability *in vivo*. These additional signals reside in the C-terminal half of Mrp17. This again indicates, that the mitochondrial targeting regions are scattered over the Mrp17 sequence, and for this protein, the N-terminal region is irrelevant for efficient mitochondrial import.

### Characterizing the features of the Mrp17 targeting and translocation signal

Next, we analyzed the unconventional internal targeting region of Mrp17 located between residues 30-60 in more detail. Standard prediction algorithms do not find an MTS-like sequence in this region (Fig. S3). The Mrp17 structure mostly contains beta-strands in this region and only a part of a helical stretch (Fig. 3A). Mrp17 is generally rich in positive charges (its pI is 10.5) and a high content of positive charges are a general feature of ribosomal proteins that interact with negatively charged mRNA. Interestingly, during evolution, the positive charges in MRPs (and particularly their lysine content) was further increased suggesting that positive charges might play a role beyond their relevance for neutralizing the negative charges of ribosomal RNA (Fig. S8A, B).

**Fig. 3.**
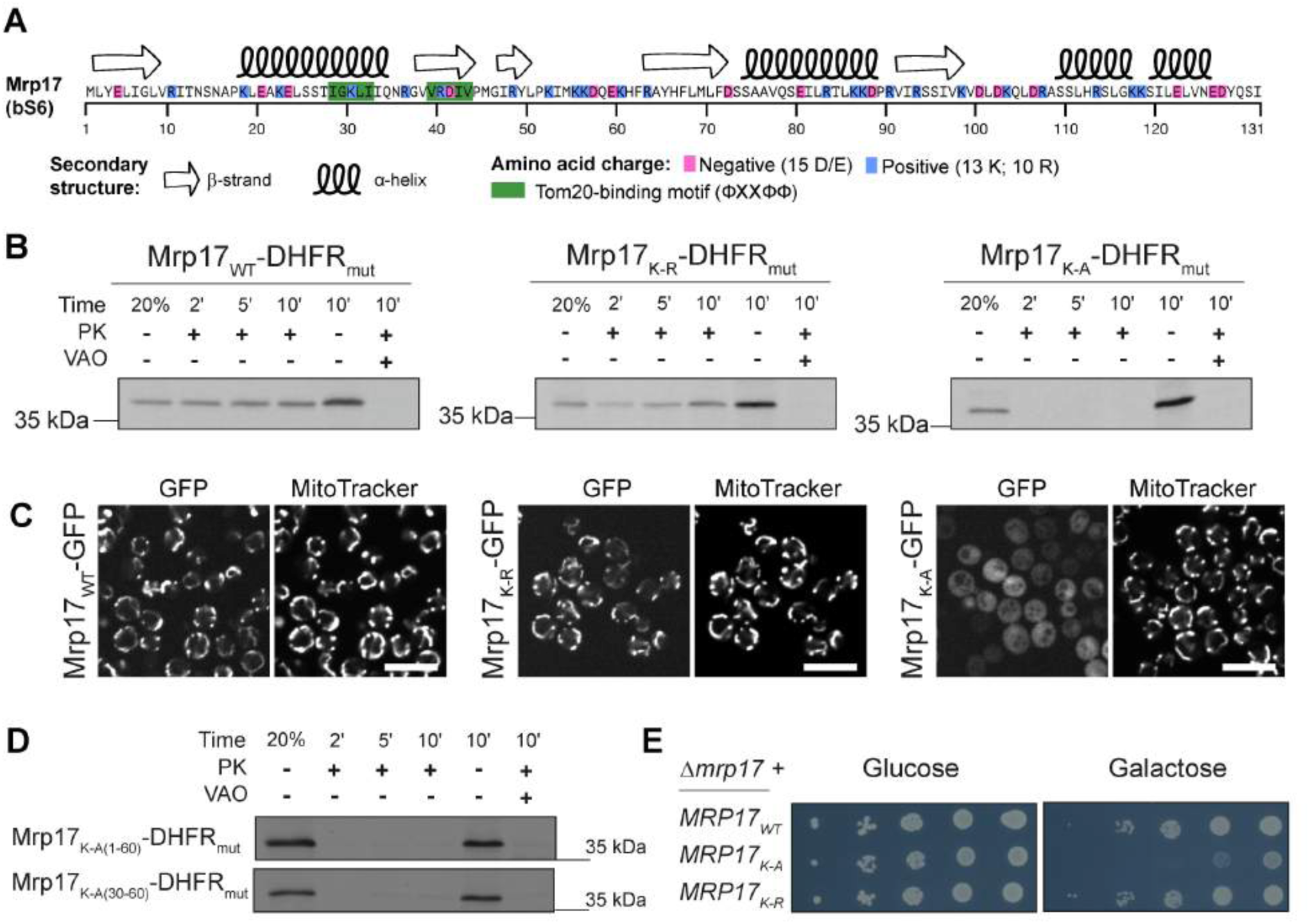
Positive charge and Tom20-binding motifs are important features of the Mrp17 targeting signal. (A) Primary sequence of Mrp17 highlighting charged amino-acids, Tom20-binding motifs and secondary structure (from PD8:5MRC); (B) In vitro mitochondria translocation capacity of WT Mrp17, Mrp17K-R with all lysines (K) substituted with arginines (R), and Mrp17K-Awith all lysines (K) substituted with alanines (A) fused to DHFRmut. Import was performed as described in Fig. 2 legend. (C) In vivo mitochondrial targeting capability of WT Mrp17, Mrp17K-R, and Mrp17K-A fused to GFP. (D) Substitution of lysines with alanines in the regions 1-60 or 30-60 of Mrp17 abolishes mitochondrial import capacity. (E) Plasmids carrying the indicated Mrp17 variants were transformed into a 6mrp17 strain that harbored an MRP17 gene on an URA3-containing plasmid. The latter plasmid was removed by plasmid shuffling and the growth of the strains was assessed on glucose and galactose media. Scale bar for all micrographs is 10 µm.

To find out whether mitochondrial targeting and import require the positive charge and if so, whether there is a specific dependence on lysines, we constructed mutants of Mrp17 with all lysines substituted with arginines thus maintaining the charge (Mrp17_K-R_) or all lysines substituted for alanines thus decreasing the charge (Mrp17_K-A_). The Mrp17_K-R_ mutant fused to DHFR_mut_ was efficiently imported into isolated mitochondria, while the Mrp17_K-A_ mutant was not (Fig. 3B). Similarly to the results obtained *in vitro*, Mrp17_K-R_ fused to GFP and expressed in yeast colocalized with mitochondria as the wild type Mrp17 while Mrp17_K-A_-GFP remained cytosolic (Fig. 3C, Fig. S8C). Thus, the positive charge is an important feature of the Mrp17 translocation signal. We also constructed Mrp17 mutants with lysines substituted with alanines only in the 1-60 or 30-60 regions. These mutants fused to DHFR_mut_ were also not imported indicating that the positive charges indeed must be positioned in the targeting signal region (Fig. 3D). Interestingly, Mrp17_K-R_ was able to rescue the growth defect of *Δmrp17* cells on respiratory media suggesting that lysines per se are not essential for import and assembly of Mrp17, but the positive charges are (Fig. 3E).

Using mutagenesis, we determined that positive charge, most critically between amino-acids 30-60, is important for the function of Mrp17 targeting and translocation signal.

### Determining the import pathway of Mrp17

Since Mrp17 lacks the typical features and targeting signals of matrix proteins, we wondered how this protein was first recognized by mitochondria and later imported. To test the receptor requirement for the mitochondrial import of Mrp17, we treated isolated yeast mitochondria with trypsin that removes all the import receptors of the Translocate of the Outer Membrane (TOM) complex from the mitochondrial surface but spares membrane-embedded TOM subunits (Ohba and Schatz, 1987). Trypsin treatment strongly inhibits the import of MTS containing matrix proteins such as Atp1 (Fig. 4A) but does not affect the import of most intra-membrane space (IMS) proteins (Gornicka et al., 2014; Lutz et al., 2003). The import of Mrp17-DHFR_mut_ was even more sensitive to trypsin treatment than the import of Atp1 (Fig. 4A). Thus, Mrp17 import into mitochondrial matrix strongly depends on the presence of the mitochondrial outer membrane receptors.

**Fig. 4.**
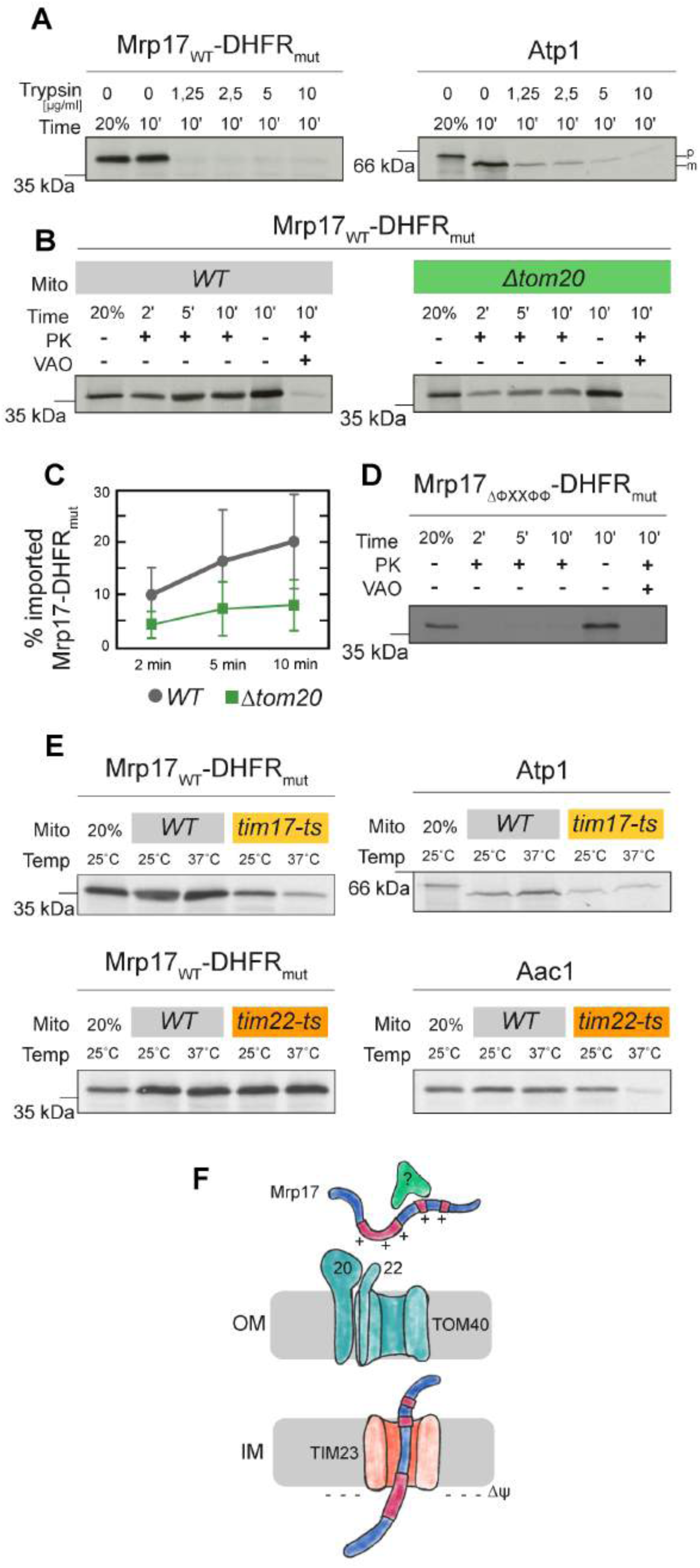
Mrp17 requires Tom20 for translocation into isolated mitochondria. (A) The in vitro import of Mrp17-DHFRmut is sensitive to the elimination of outer mitochondrial membrane proteins by trypsinization (left), more so than the model import substrate Atp1 (right). Mitochondria were incubated with the indicated concentrations of trypsin and the import assay was performed as described in the legend for Fig. 2. Proteinase K (PK) was not added to the sample without mitochondria. (8) In vitro translocation of Mrp17-DHFRmut into mitochondria isolated from I/VT and t:,,tom20 yeast showing reduced translocation in *t:,,* tom20 background. (C) Quantification of the experiment in panel (8) repeated 3 times, whiskers correspond to minimum and maximum values. (D) Substitution of both Tom20-binding motifs (ФXXФФ) with alanines abolishes Mrp17 mitochondrial import capacity. (E) Import of Mrp17-DHFRmut and control proteins Atp1 and Aac2 into I/VT, tim17-ts and tim22-ts mitochondria, import was performed at indicated temperatures, proteinase K was added to all samples except the loading control in the first lane of each autoradiograph (20% of protein amount used for each import reaction). (F) Mrp17 (blue) targeting and translocation model: positively charged internal signals (magenta) might bind yet unidentified targeting factors (green) and then are recognized at the outer membrane of mitochondria by the same receptors Tom20 and Tom22 as a regular MTS, then imported through TOM40 (teal) and TIM23 (orange) translocons.

To determine the dependence of Mrp17 import on individual receptors we purified mitochondria from yeast lacking Tom20 (Δ*tom20*), the main receptor for recognition of canonical MTSs. Mrp17-DHFR_mut_ import into Δ*tom20* mitochondria was strongly reduced (Fig. 4B,C) indicating that Tom20 is involved in the import process. Deletion of Tom70 and its lowly expressed paralog Tom71 had no effect on the import of Mrp17-DHFR_mut_ (Fig. S9), consistent with the absence of internal MTLS-like sequences (iMTS-Ls, (Backes et al., 2018)) in the protein (Fig. S3).

Since Tom20 deletion can affect the import of many mitochondrial proteins, its effect on Mrp17 can be indirect. However, we could find two potential motifs for binding Tom20 (ФXXФФ, where Ф is hydrophobic amino acid and X - any amino acid; Fig. 3A)(Muto et al., 2001; Obita et al., 2003). To test whether the requirement for Tom20 binding is direct we mutated the Tom20-binding motifs in the Mrp17 translocation signal. For this, we constructed a mutant where each amino acid of both Tom20 binding motifs was substituted with alanine - Mrp17_ΔФXXФФ_. Indeed, the mutant fused to DHFR_mut_ was not imported into isolated mitochondria supporting an essential role of these Tom20-binding motifs for import (Fig. 4D). However, the residual translocation of Mrp17-DHFR_mut_ in Δ*tom20* compared to trypsinized mitochondria suggests that Mrp17 can use additional receptors, albeit at lesser efficiency. Such a bypass receptor could potentially be Tom22 which was shown to work in concert with Tom20 in MTS recognition (Yamano et al., 2008).

Next, we used the temperature-sensitive mutants *tim17-ts* and *tim22-ts* that reduce the protein import along either of the two Translocase of the Inner Membrane (TIM) pathways - the TIM23 and TIM22 pathways, respectively. Mrp17 import into *tim17-ts* mitochondria was reduced whereas its import into *tim22-ts* mitochondria was not affected showing that Mrp17 uses the TIM23 pathway, similar to canonical MTS-containing proteins (Fig. 4E).

To summarize, Mrp17 has an internal targeting signal that shares some properties such as positive charge with a regular MTS. Indeed, its import pathway is similar to MTS containing proteins suggesting that mitochondrial components recognize it as a *bona fide* MTS yet even state-of-the-art prediction algorithms do not, suggesting that we lack information on certain MTS characteristics (Fig. 4F).

### Comparison of Mrp17 to its bacterial homolog

Unlike many other core MRPs that significantly extended their structures with insertions and N/C-terminal expansions, Mrp17 maintained the overall structure of its bacterial ancestors (Fig. 5A, Fig. S10). This means that in the course of evolution, the acquisition of an N-terminal MTS was either not possible so that the Mrp17 mitochondrial targeting signal had to be accommodated within the existing “bacterial” structure, or the ribosomal protein was already predisposed for mitochondrial import. Since RNA-binding regions show similarity to mitochondrial targeting signals with both requiring positively charged and hydrophobic amino acids - this was an appealing hypothesis.

**Fig. 5.**
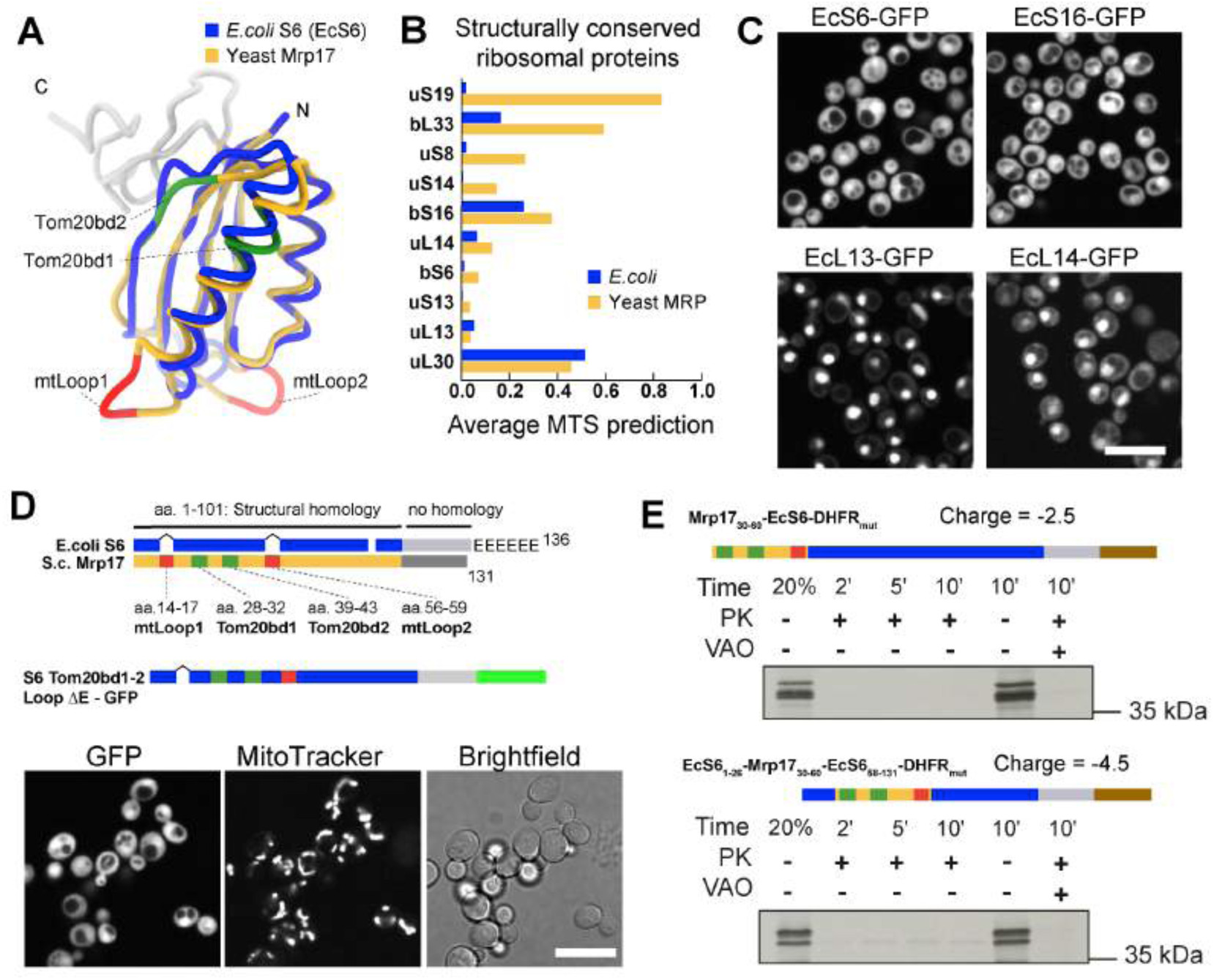
Bacterial homologs of MRPs have no mitochondrial import predisposition. (A) Structural alignment of the yeast Mrp17 (from PDB:5MRC) and its E. coli homolog ribosomal protein S6 (from PDB: 6WDO) viewed from its cytosol-facing side, highlighting regions of structural homology (blue and yellow) and non-homologous C-terminal regions (grey), Tom20-binding domains in Mrp17 are highlighted in green and expanded loops in red, schematic summaries of such structural alignments for all MRPs and their bacterial homologs are shown in Fig. S10. (B) Average MTS prediction scores (MitoFates and TargetP2) for structurally conserved yeast MRPs and their bacterial homologs sorted by the difference between bacteria and mitochondria. (C) Expression of structurally conserved bacterial RPs fused to GFP in yeast cells. (D) Schematic summary of structural alignment of bacterial S6 and yeast Mrp17 shown in panel (A) highlighting additional features of Mrp17 structure (two loops and two Tom20-binding sites) and six glutamate residues (E’s) at the C-terminus of bacterial S6 (top), schematic of chimeric construct of EcS6 that lacks C-terminal glutamines, but incorporates structural features of Mrp17 and is fused to GFP shown bright green (middle), and the result of expression of this chimeric construct in yeast (bottom). (E) Chimeric variants of EcS6-DHFRmut with Mrp17 amino acids 30-60 fused to it at the N-terminus or incorporated instead of the homologous region of S6 itself (top), and in vitro import of these constructs into isolated yeast mitochondria. Lanes are labeled as described in Fig. 2 legend (bottom). Scale bar in all micrographs is 10 µm.

To investigate both possibilities, we compared average MTS prediction scores of Mrp17 (and other structurally conserved MRPs) to their closest bacterial homologs with known structure, i. e. *Escherichia coli* ribosomal proteins (Fig. 5B, Fig. S10). Two out of the 10 structurally conserved proteins that lacked cleavable MTS, indeed had much higher MTS prediction scores in mitoribosomes compared to bacterial ribosome meaning that uncleavable MTS-like signals were developed at their N-termini. For other proteins, both homologs had equally low scores (less than 0.5) indicating that some sequence properties of bacterial and mitoribosomal proteins are similar.

Next, we tested if these bacterial proteins already have properties that allow them to be imported into mitochondria *in vivo* by expressing them as GFP fusions in yeast (Fig. 5C, Fig. S11A). None of the proteins localized to mitochondria showing that these BRPs do not have an intrinsic mitochondrial targeting capacity and that targeting signals were incorporated into conserved MRP structures in the course of evolution. Interestingly, two of the proteins (EcL13 and EcL14) had an intrinsic nuclear targeting capacity which was tolerated by the cells (Fig. S11B).

We aimed to check if any particular sequence or structural features present in Mrp17 compared to its *E. coli* homolog, protein S6 (EcS6) can be responsible for mitochondrial targeting. Most prominent of these features are two loop extensions (the first one is not in the range of amino acids 30-60 but the second is at the end of it and contains additional positive charges) and the two Tom20-binding motifs (Fig. 5A,D). We therefore introduced these four Mrp17 features into the homologous regions of EcS6 altogether or in different combinations, but none supported mitochondrial targeting of the chimeric constructs fused to GFP (Fig 5D, Fig. S11C). *In vitro*, adding to EcS6 the whole targeting sequence of Mrp17 (from amino acids 30 to 60, sufficient for translocation of DHFR_mut_) was still not sufficient for translocation of EcS6-DHFR_mut_ when fused to it at the N-terminus (Fig. 5E) suggesting that the overall negative charge of EcS6 (−6.5, without C-terminal glutamines) strongly inhibits mitochondrial import of chimeric constructs. When the same domain, Mrp17_30-60_, was introduced into its homologous position in EcS6 and fused to DHFR_mut_, such chimeric construct showed some import capacity (Fig. 5E). This indicates that in addition to the charge effects, the import signal contained in Mrp17_30-60_ is also dependent on its spatial context to be functional.

Based on the comparison with the bacterial homolog, we suggest that the main features ensuring mitochondrial import of Mrp17 are the Tom20-binding motifs and its overall positive charge both of which were acquired without significant structural rearrangements.

## Discussion

In this work we collected the available information on the MTSs of yeast MRPs from proteomic and structural studies as well as MTS prediction algorithms. Using *in vivo* structure-function assays we found that some MRPs possessed internal targeting signals that can be poorly predicted. We characterized the internal targeting signal of Mrp17 in more detail and found that it shares some features and import pathway with regular MTS-containing proteins.

Why would Mrp17 and other MRPs rely on internal signals instead of evolving a canonical N-terminal MTS? For Mrp17 and other MRPs that originated from bacterial ancestors and have not acquired any additional sequence extensions - they may have been simply because they did not need to evolve new signals as they already possessed mitochondrial import capacity due to their positive charge and hydrophobicity that are needed to bind ribosomal RNA. Such intrinsic import capacity was indeed shown for some *E. coli* proteins (Lucattini et al., 2004). However, in our work even strongly positively charged *E. coli* ribosomal proteins S12, S16, L13, and L14, although structurally conserved with their MRP homologs, were not targeted to mitochondria when expressed in yeast suggesting no initial targeting predisposition (Fig. 5C). Instead, we propose that the N-termini of these proteins were too conserved to be extended with a cleavable MTS. So proteins, such as Mrp17, had to accommodate a targeting signal elsewhere in their sequences.

The N-terminus of Mrp17 is tightly positioned on the contact with neighboring proteins (Fig. 6A, Fig. S12A). If this interface arose before the final maturation of the mitochondrial protein import system, it would create challenges for the evolution of an N-terminal targeting signal. In principle, such and engagement of the N-terminus in a structural interface can still be kept with a cleavable MTS if proteases cleave the protein precisely before the first structurally conserved residue. Indeed, some MRPs have evolved in that way (Fig. S10). We hypothesize that for Mrp17, however, such precise cleavage could not evolve. First reason for that is that the two first amino acids were too conserved to be adjusted for a cleavage site. Second, an intermediate stage of cleavable MTS evolution, which could be a sub-optimally cleaved N-terminus with few extra amino acids, could not be accommodated in the structure, so it never got to evolve a precisely cleaved MTS. Instead, Mrp17 remodeled the charge for the whole protein and developed internal targeting signals. Interestingly, the main signal identified in our work was not positioned in structurally distinct areas that are different between Mrp17 and EcS6 (Fig. 5A,D; loop expansions, C-terminal region) but mostly distributed between amino acids 30-60 which form a lot of new protein-protein contacts within the mitoribosome compared to the bacterial ribosome (Fig. S12A,B). So, the Mrp17 targeting signal might have co-evolved with protein-protein interactions which is similar to the evolution of nucleolar-targeting signals in cytosolic ribosomal proteins (Melnikov et al., 2015).

**Fig. 6.**
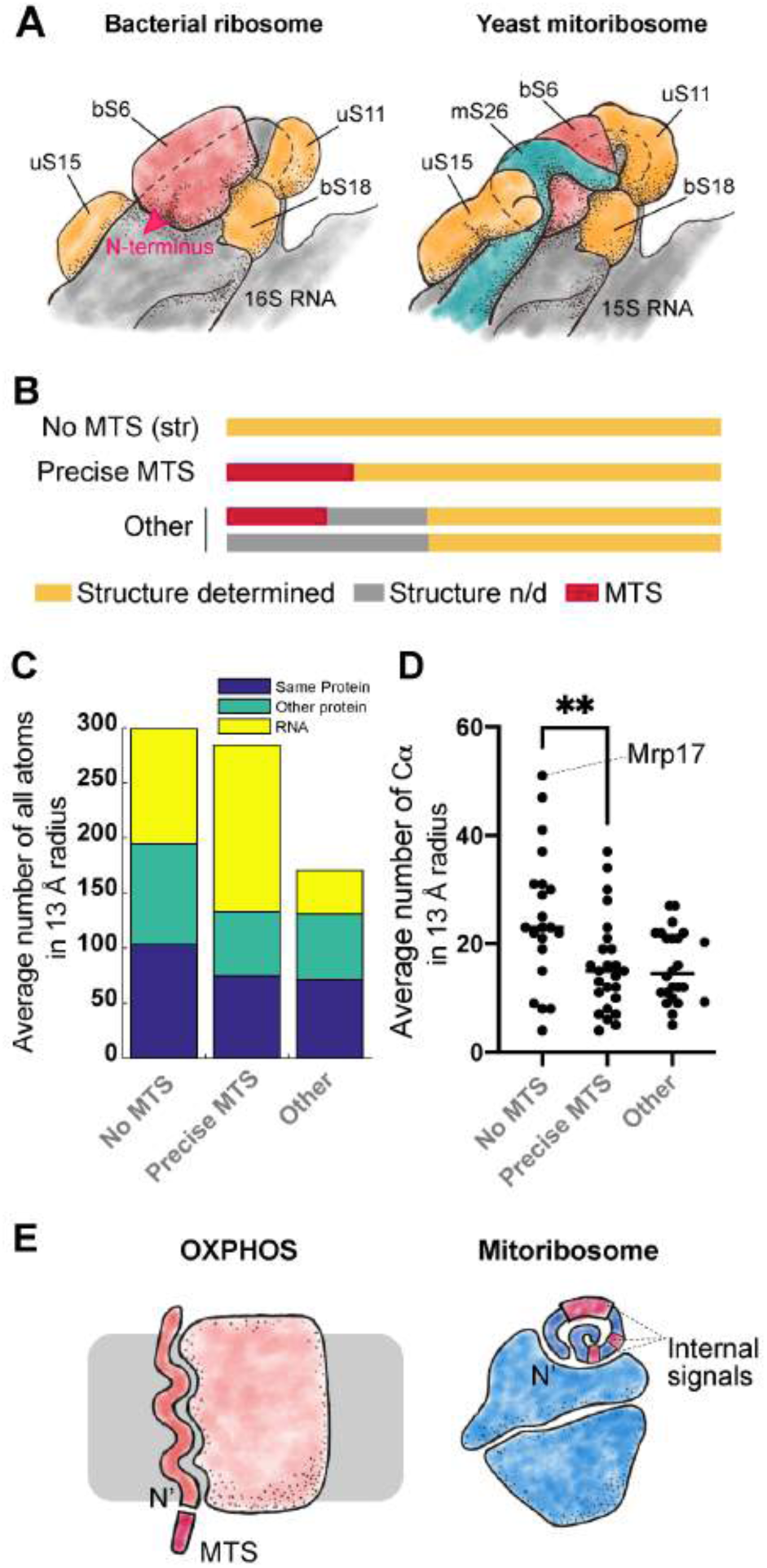
Structural constraints for the evolution of N-terminal mitochondrial targeting signals. (A) Schematic depiction of the S6 protein (red) in the context of bacterial ribosome (after PDB:6WD0, left) and yeast mitochondrial ribosome (after PD8:5MRC, right) highlighting the position of the N-terminus of S6 (red arrow, only shown on the left image). Proteins homologous between bacterial and yeast structures are shown in orange, mitochondria-specific protein mS26 in teal and rRNA in grey. (B) MRPs divided in three groups depending on relative position of the MTS and structured residues: No MTS (str) - structure starts at the very N-terminus without cleavable MTS, Precise MTS - MTS is cleaved off 1-2 amino acids before the structure starts, Other - unstructured uncleaved amino acids present before the structure start. (C) MRP N-termini solvent exposure: average number of atoms belonging to the same protein, other protein chains or RNA in the 13A radius from the first structured amino acid C atom, plotted separately for proteins with their most N-terminus appearing in the structure vs proteins with precisely cleaved MTS (first structured amino acid is within 3 or less amino acids from the end of the MTS) and other proteins which have a longer unstructured extension calculated based on PD8 : 5MRC. (D) Same as (E) but only coordination number (number of Cl:iil atoms in the 13A radius (Hamelryck, 2005)) is calculated. (F) Compared to OXPHOS complex components that usually have their N-termini free to develop an MTS (left), mitoribosomes often employ the MRP N-termini in binding interfaces which puts structural constraints on the development of N-terminal MTS and promotes the evolution of internal targeting sequences that can also take part in ribosome assembly (right).

Can structural constraints also explain why other MRPs did not develop cleavable MTSs? To assay that, we divided all MRPs into three groups according to the relative position of their MTS cleavage site (if present) and the first structured residue (Fig. 6B). We hypothesized that despite being involved in structure formation, the proteins that developed an MTS precisely cleaved off before the first structured residue might have N-termini less buried in the structure that could provide some flexibility for MTS cleavage evolution compared to the proteins that failed to develop an MTS and have a deeply buried N-terminus, like Mrp17. However, this was not the case and these two groups had their N-termini equally deeply buried in the structure (Fig. 6C, “No MTS” and “Precise MTS” groups). The validity of such measurement was confirmed by the fact that the MRP N-termini with few more unstructured residues before the structure start are indeed more often positioned on the ribosome surface (Fig. 6C, “Other” group). Thus, there is no strict limitation for a buried N-terminus to develop an MTS.

Interestingly, we found that the N-termini without a cleavable MTS had more protein and less RNA around them in the ribosome structure with Mrp17 being an extreme example (Fig. 6B,C). This suggests that unlike protein-protein contacts, protein-RNA contacts can be more permissible to the change in the protein component that will allow the adjustment of the N-terminus for protease cleavage. Our observation agrees with the study of sequence conservation of ribosomal proteins that revealed that amino acids on the protein-RNA interfaces are indeed less conserved than the amino acids on the protein-protein interfaces (Pilla and Bahadur, 2019). Supposedly, RNA-protein interfaces are the most ancient and important for ribosome assembly, so their smaller conservation compared to protein-protein interfaces might seem counterintuitive (Fox, 2010). However, this could be explained by the nature of protein-RNA binding that often relies on positively charged and hydrophobic amino-acids while the fine shape of the protein surface is not so important. On the other hand, protein-protein interfaces very much rely on the exact match of surface shapes which makes amino-acid substitutions much more detrimental for the interface integrity.

Still, involvement in protein-protein contacts does not fully explain the absence of a cleavable MTS in some less deeply buried MRPs (Fig. 6C) indicating that other factors might be at play. Some possible explanations can be that a new MRP is recruited to the ribosome with its targeting signal already positioned elsewhere in its sequence. Another option is that a recruited protein has a normal cleavable MTS but after recruitment, the MTS loses its cleavage site and becomes an integral part of the structure still positioned on the ribosome surface. It is also possible that in some MRPs the flexible N-terminal extensions play other important roles incompatible with N-terminal targeting signal properties. In the future, more systematic analysis of mitoribosome structures from different organisms can shed light on the complex interplay between the evolution of mitoribosome assembly on one side and the targeting signals of its components on the other.

To summarize, we suggest that at least some of the non-canonical targeting signals can develop in MRPs not only to fulfill certain functions, such as in the case of Mrp10 and Mrpl32 (Bonn et al., 2011; Longen et al., 2014), but under the pressure of structural constraints imposed by the mitoribosome assembly. It seems that unlike OXPHOS complexes, which also underwent complex evolution in mitochondria (Sluis et al., 2015), protein components of mitoribosome more often use their N-termini to establish important interactions and thus could not so easily develop a cleavable MTS (Fig. 6D). Instead, they developed multiple internal targeting signals of different strengths such as those found in Mrp17. This might have been relatively easy for ribosomal proteins that either already contain a lot of positive charges, or can easily increase their content due to abundant protein-RNA contacts. An alternative strategy would be to place a targeting signal on the C-terminus. For now, the only known matrix protein with a C-terminal targeting signal is Hmi1 that also cannot tolerate an N-terminal extension (Lee et al., 1999). Such diversity of mitochondrial targeting signals highlights the need for better targeting signal prediction algorithms.

In this work we have shown that many MRPs have unconventional, non-cleavable targeting signals that can use the same import pathway as a regular MTS. This demonstrates an incredible versatility of the mitochondrial import system that can accommodate such a range of substrates and poses the question what is the mechanistic basic of balancing such versatility with the specificity of protein import process.

## Acknowledgements

We are grateful to Prof. Ada Yonath and Dr. Anat Bashan for their valuable comments and discussion, and to Victor Tobiasson and Alexey Amunts for their advice on structure analysis. We thank Amir Fadel for technical help with microscopy and growth assays, and Nikolaus Pfanner for temperature-sensitive yeast mutants. We thank Einat Zalckvar for critical reading of the manuscript.

Collaborative work between Herrmann and Schuldiner labs is supported by a DIP collaborative grant (MitoBalance). Work in the Schuldiner lab is supported by an ERC CoG (OnTarget 864068). We are grateful for funding from the Deutsche Forschungsgemeinschaft (HE2803/9-1 to J.M.H.). Y.B. is supported by EMBO Long-term postdoctoral fellowship. M.S in an incumbent of the Dr. Gilbert Omenn and Martha Darling Professorial Chair in Molecular Genetics. The robotic system of the Schuldiner lab was purchased through the kind support of the Blythe Brenden-Mann Foundation.

## Materials and Methods

### Yeast strains and plasmids

All yeast strains used for fluorescent protein expression and imaging were constructed on the BY4741 haploid or BY4743 diploid background (Brachmann et al., 1998). For mitochondria purification we used W303 background and for growth rescue with plasmid shuffling - YPH499 background. The strains used in this study are listed in Table S2. All strains were constructed using standard LiAc/ssDNA/PEG based transformation protocol (Gietz and Woods, 2006). For transformation we used the standard plasmids for PCR-based tagging and knock outs (Janke et al., 2004; Longtine et al., 1998) and plasmids generated in this study (Table S3). Mutant versions of Mrp17 genes and genes for bacterial ribosomal proteins optimized for expression in yeast were ordered from GeneWiz or Genescript. When strains were constructed by genomic integration, primers were designed using Primers4Yeast (http://wws.weizmann.ac.il/Primers-4-Yeast)(Yofe and Schuldiner, 2014) or manually if the gene was truncated. Primers used in this study are listed in Table S4.

### Yeast growth

Yeast cells were grown on either liquid media or solid media that contained 2.2% agar. For selection only for antibiotic resistance yeast cells were grown on YPD media (2% peptone, 1% yeast extract, 2% glucose) supplemented with nourseothricin (NAT) to 0.2 g/l. For auxotrophic selections yeast were grown in SD media (0.67% yeast nitrogen base without amino acids and with ammonium sulfate, 2% glucose, and OMM amino acid mix (Hanscho et al., 2012)) if necessary supplemented with the same amount of antibiotic.

### Growth assays

For growth assays with plate reader the strains were inoculated in 96-well plate 1 day before the experiment start if grown in fermentative media (YPD, see above) or 2 days before the experiment if grown in fermentative media YPGlycerol (2% peptone, 1% yeast extract, 2% glycerol) at 30°C with 500 rpm shaking in automated Liconic incubator. On the day of experiment the saturated cultures were diluted 1:50 in fresh media using EVO Freedom liquid handler (Tecan) and incubated at 30°C with shaking. The optical density measurements at 600 nm were taken every 30 min with the SPARK plate reader (Tecan). Each assay was repeated 2 times (Fig. S5).

For drop dilution assay, the yeast cultures in the phase of exponential growth (OD_600_ ∼ 0.6) were sedimented and diluted in fresh media to the OD=0.1. This suspension was serially diluted *×*10 and 2.5 µl of each dilution was spotted on the agar plate using a multichannel pipette. The plates were incubated for 2 days at 24°, 30°, and 37°C and imaged using smartphone camera.

## Analysis of translation start site using ribosome profiling

To check for possible mis-annotation of the translation start sites of MRPs in the *Saccharomyces* genome database, we re-analyzed data from a ribosome profiling study that was specifically designed to detect translation initiation sites (ref. Knöringer, Groh, Krämer et al., in preparation; data available at GEO with the accession number GSE172017, access code available upon request). Briefly, ribosome profiling libraries of yeast cells (YPH499) were prepared as described (Stein et al., 2019) with the following modification: In one replicate, 100 µg/ml cycloheximide was added to the yeast culture 2 min before harvesting and lysis, while in the other replicate, cells were not in contact with cycloheximide prior to cell lysis. Cycloheximide inhibits translation elongation, but not translation initiation, which results in an enrichment of ribosome footprints at translation initiation sites in CHX-treated samples compared to untreated samples. Ribosome footprints were sequenced and aligned to the yeast genome. Footprint densities along the annotated open reading frames of MRP genes were analyzed and the translation start site was re-annotated based on the following criteria: (1) the presence of contiguous ribosome footprints in all samples (previously annotated introns were taken into consideration); (2) the presence of an enrichment of footprints in the CHX-treated sample at the very 5’ position of the suspected ORF; (3) the presence of an AUG codon at the suspected start site.

### Fluorescence microscopy

The day before the experiment yeast were grown to saturation. On the day of experiment the saturated culture was diluted 1:50 in SD media without selection or with just auxotrophic selections. Cells were grown for 4 h and applied on glass-bottom plates coated with concanavalin A and left for 20 min to adhere. If mitochondria needed to be visualized, the solution was removed from the wells and 50 nM MitoTracker Orange CMTMRos (ThermoFisher #M7510) diluted in imaging media (SD media with complete set of amino acids but without riboflavin) was placed in the wells for 10 min. Imaging was performed in fresh imaging media. Cells were imaged using VisiScope Confocal Cell Explorer system consisting of Olympus IX83 microscope, Zeiss Yokogawa spinning disk scanning unit equipped with PCO-Edge sCMOS camera controlled by VisiView sofware. Images were recorded with 488 nm laser illumination for GFP channel and 561 nm laser illumination for MitoTracker Orange and 60*×* oil objective. Micrographs were cropped, and slightly adjusted for brightness and contrast using Fiji (Schindelin et al., 2012).

### Data analysis

The data on the first amino acid in the mitoribosome structures were extracted directly from the mmCIF files of PDB entries 6WD0 (Loveland et al., 2020), 5MRC (Desai et al., 2017), 6YWS, 6YW5 (Itoh et al., 2020), 6NU2 (Koripella et al., 2019), 6GAW (Kummer et al., 2018), 6XYW (Waltz et al., 2020), 6HIV (Ramrath et al., 2018), 6ZP1 (Tobiasson and Amunts, 2020) using PDBeCIF (https://pypi.org/project/PDBeCif/). The number of atoms around each C-alpha atom were calculated from this data using Python (McKinney, 2010).

The cleavage site in MRP N-termini was annotated from the following sources with priority as listed: (1) original publication with N-terminal sequencing as cited by UniProt (Boguta et al., 1992; Dang and Ellis, 1990; Davis et al., 1992; Graack et al., 1988; Graack et al., 1991; Grohmann et al., 1989; Grohmann et al., 1991; Kitakawa et al., 1990; Kitakawa et al., 1997; Matsushita and Isono, 1993; Matsushita et al., 1989), (2) high-throughput N-terminal proteomic dataset (Vögtle et al., 2009), (3) cleavage site prediction by UniProt, (4) cleavage cite prediction using MitoFates (Fukasawa et al., 2015). If the cleavage site annotation didn’t agree with the structural data (cleavage annotated after the amino acid actually present in the structure), we took the annotation from the source of next priority. If none of the annotations agreed with the structure, the N-terminus cleavage site was marked as “NA” (Table S1).

Mitochondrial targeting sequence prediction scores were calculated using MitoProt (Claros and Vincens, 1996), TargetP1 (Emanuelsson et al., 2000), TargetP2 (Armenteros et al., 2019), and MitoFates (Fukasawa et al., 2015) as described. The profiles and propensities of iMTSLs were calculated as described (Backes et al., 2018; Boos et al., 2018). Protein charge and hydrophobic moments were calculated using EMBOSS suite (Rice, 2000).

Structures were visualized using ChimeraX (Goddard et al., 2018), and electrostatic potentials using Chimera (Pettersen et al., 2004). Structural alignments were performed using FATCAT (Li et al., 2020), sequence alignment was performed and visualized using UniPro UGENE (Okonechnikov et al., 2012).

All other data analysis was performed using MATLAB (MathWorks). Plots were produced using MATLAB, Microsoft Excel, R (R Core Team, 2020), and GraphPad Prism.

### Miscellaneous

The following procedures were carried out as published before: isolation of mitochondria and *in vitro* import experiments (Peleh et al., 2015) and plasmid shuffling (Backes et al., 2019).

List of Supplementary Materials

Figs. S1-S12

**Table S1.** Manual annotation of cleavable MTS’s and other properties of yeast MRPs, Legend appears in Table.

**Table S2.** *In vivo* characterization of 15 MRP N-termini targeting capacities. Legend appears in table.

**Table S3.** Yeast strains used in this study.

**Table S4.** Plasmids used in this study.

**Table S5.** Primers for yeast transformation used in this study.

---

**Fig. S1.**
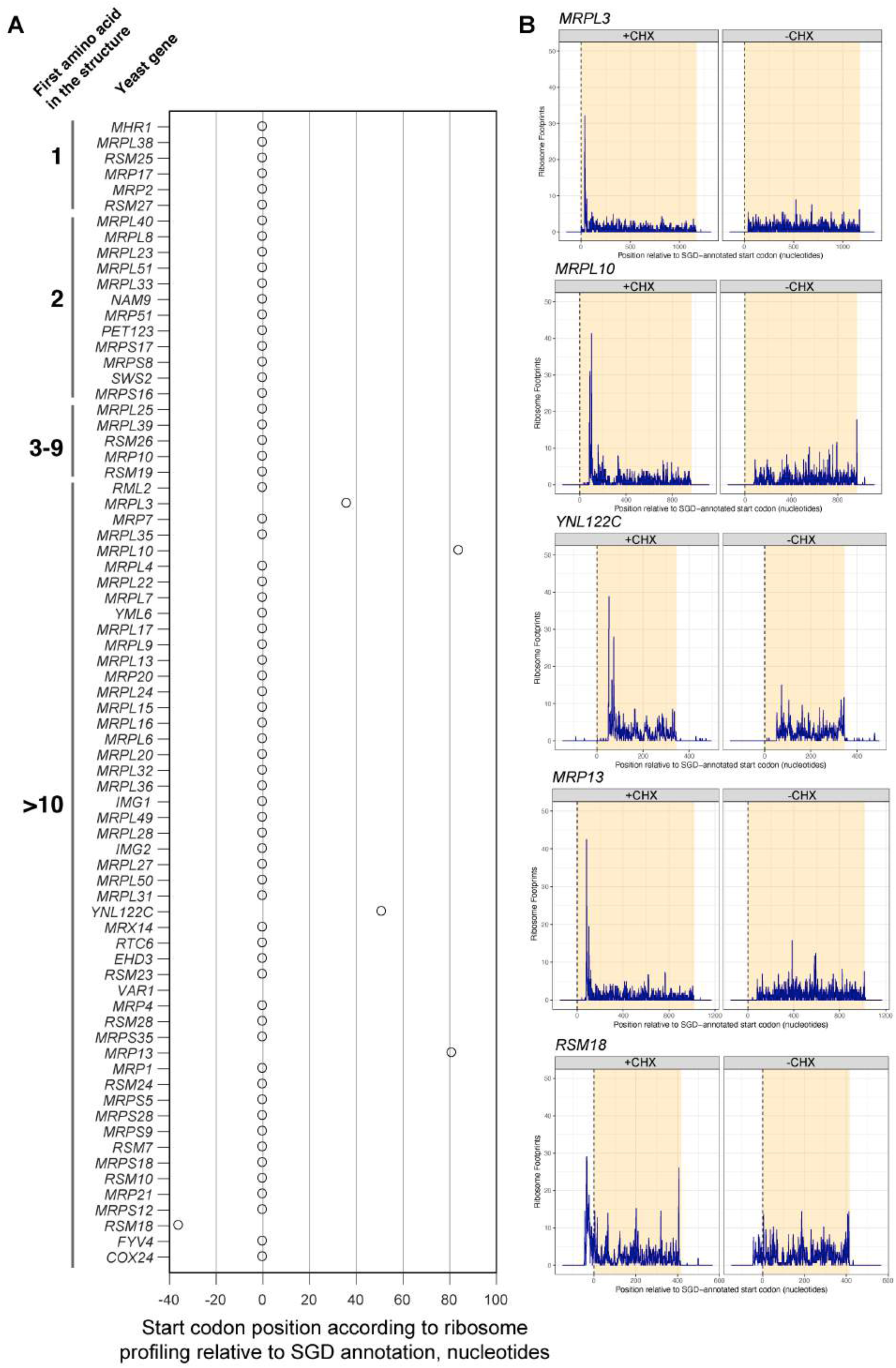
RiboSeq reveals no mis-annotated upstream translation starts thatcan account for the missing N-terminal signals. (A) For each MRP the translation start measured by RiboSeq is plotted relative to SGD annotation, revealing that only one protein has an N-terminal extension while 4 others have shorter N-termini, all of the misannotated proteins are reported to have an MTS (Table S1); no MRPs with their N-termini in the structure (first structured amino acid <10) had misannotated translation starts. (8) Ribosome footprints for each ORF with misannotated start codons in the presence of cycloheximide (+CHX, left) and without cycloheximide (-CHX, right), plotted relative to the ORF annotation in SGD (shaded in yellow).

**Fig. S2.**
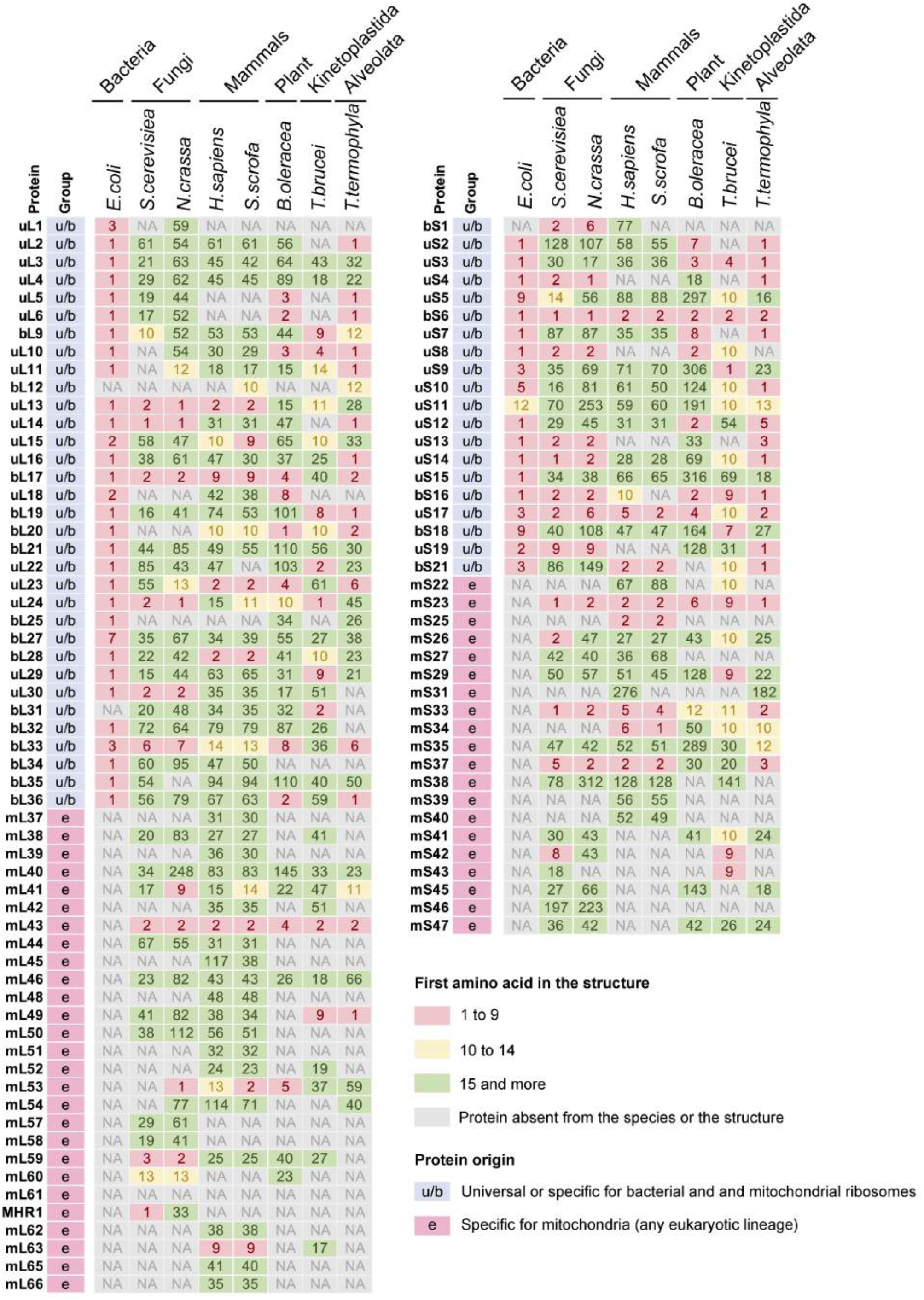
The first amino acids of MRPs found in structures of different mitoribosomes are moderately conserved. The first amino acid numbers were extracted directly from the PDB entries 6WD0 (Escherichia coli), 5MRC (Saccharomyces cerevisiae), 6YWS (large subunit Neurospora crassa), 6YW5 (small subunit N. crassa), 6NU2 (Homo sapiens), 6GAW (Sus scrota), 6XYW (Brassica oleracea), 6HIV (Trypanosoma brucei), 6ZP1 (Tetrahymena thermophila). Only MRPs occurring in two or more structures are shown.

**Fig. S3.**
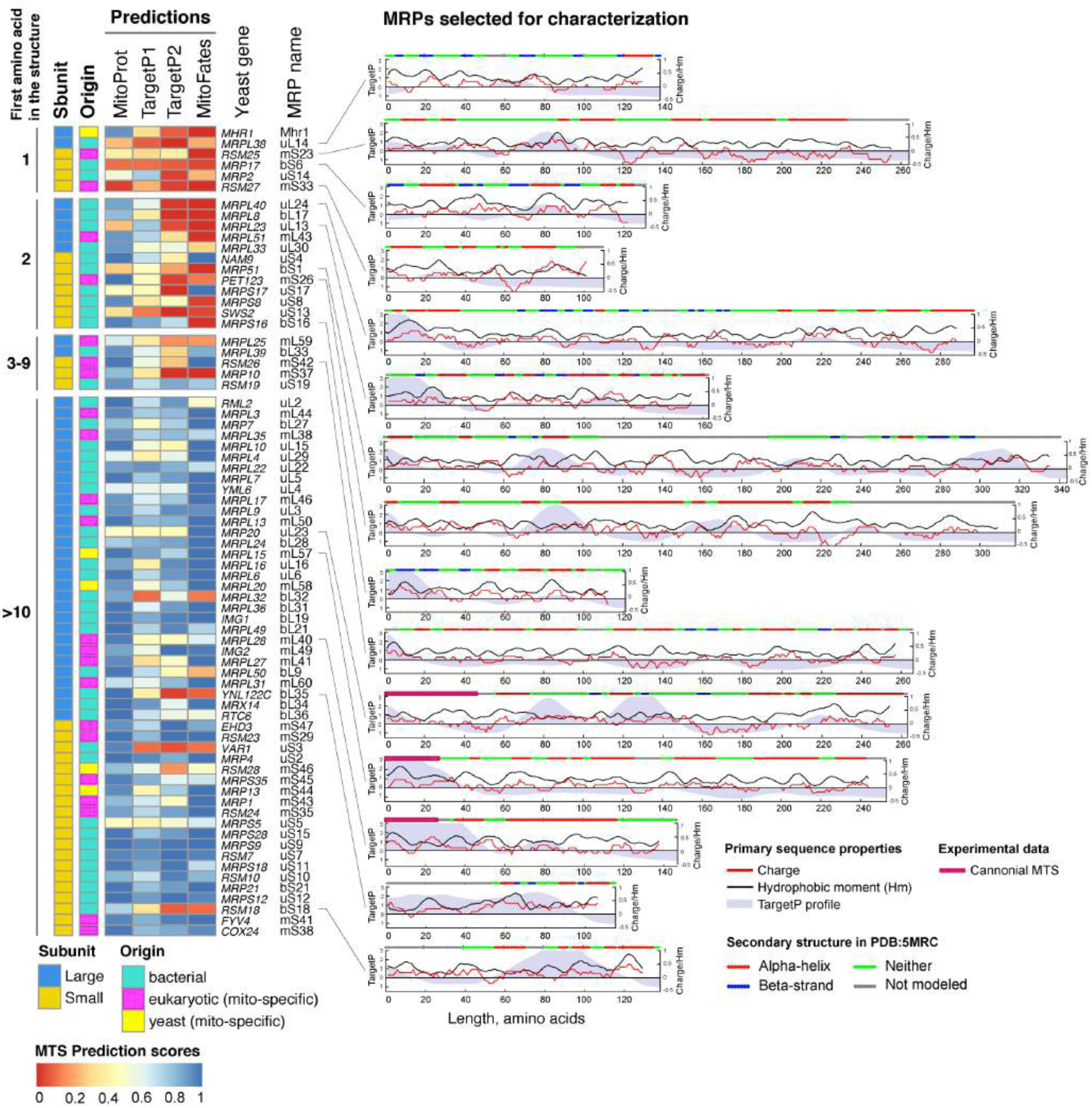
MRPs can be classified according to the presence of their most N-terminus inside the mitoribosome structure. Left - MTS prediction scores for yeast MRPs first sorted in groups by the first amino acid with reported atomic coordinates in the structure PD8:5MRC, then by subunit and then by length with protein origin and subunit noted for each MRP. Right - primary and secondary sequence properties for 15 MRPs selected for further characterization showing a variety of N-terminal and internal targeting signal predictions, overall positive charge, presence of documented cleavable MTS, and a variety of N-terminal secondary structures. Universal ribosomal protein nomenclature is used (Ban et al. 2014), except for Mhr1 which is a yeast-specific MRP.

**Fig. S4.**
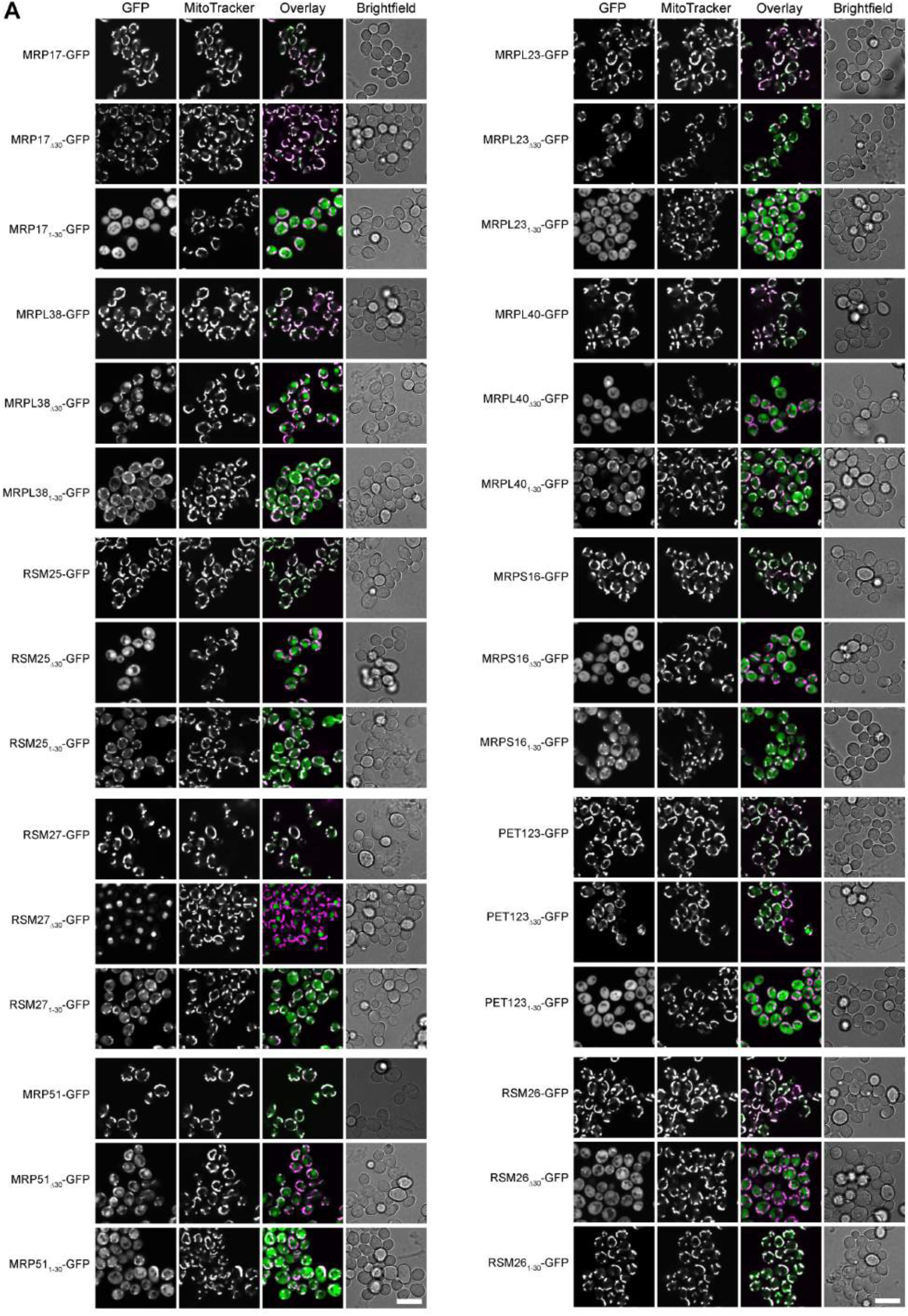
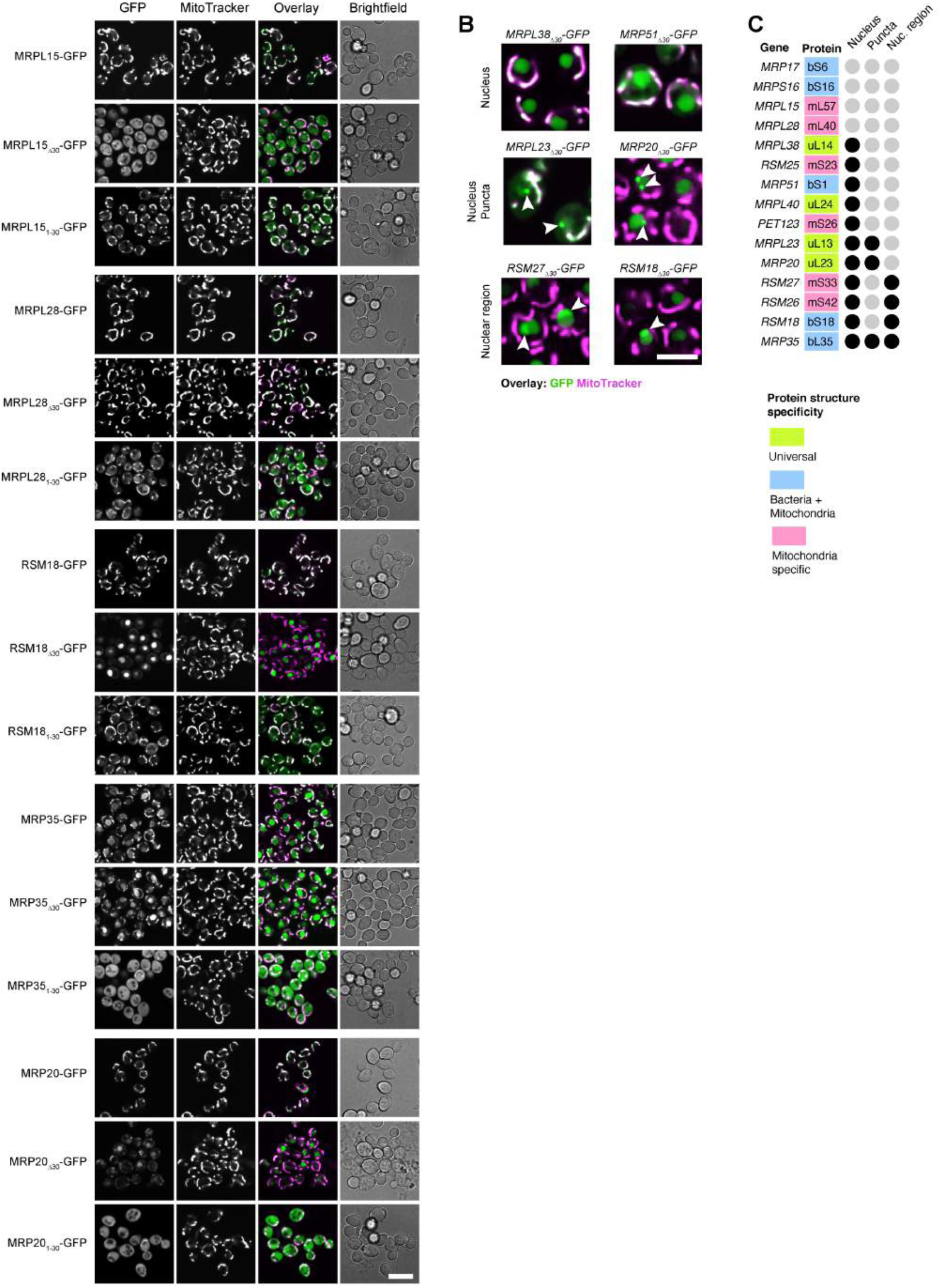
MRPs have N-termini with various targeting properties. (A) Same micrographs as in Fig. 1C shown in different fluorescent channels for all the MRP. (8) Examples of mistargeting destinations of some MRPs with deleted 1-30 amino acids from their N-termini: top row - uniform nuclear signal, middle row - nuclear signal with bright perinuclear puncta, bottom row - nuclear signal with GFP enriched in non-punctate nuclear region, shown as enlarged overlays of GFP (green) and MitoTracker (magenta) channels cropped from micrographs in panel (A). (C) Summary of mistargeting locations for one or more truncations of each MRP, if the location is observed for any of the MRP truncation, it is marked with black circle. Scale bars are 10 µm in (A) and 5 µm in (B).

**Fig. S5.**
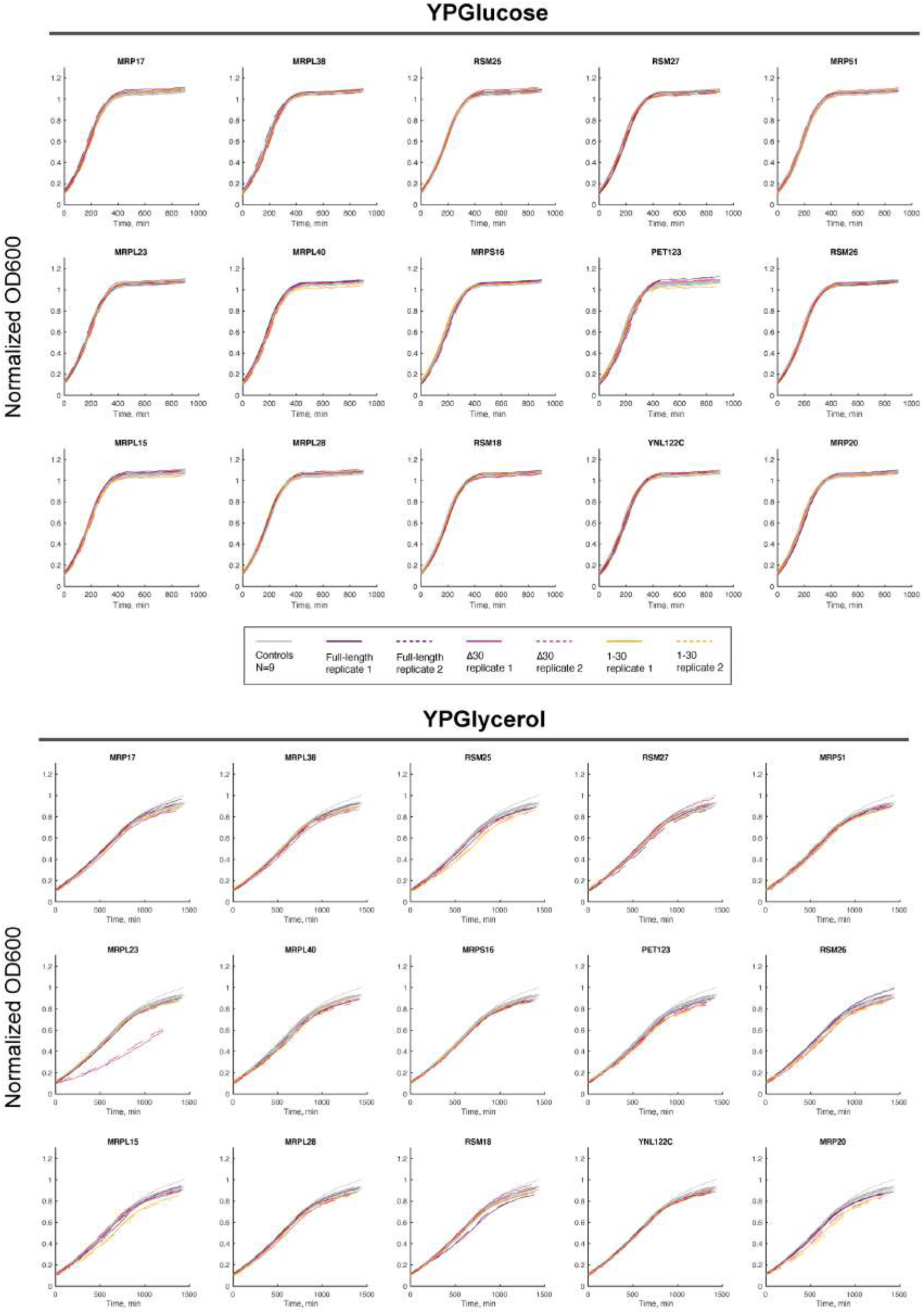
MRP truncations in the genomic locus of one allele in diploid yeast strains have no dominant negative growth defects in fermentative media and rare defects in respiratory media despite having non-mitochondrial localizations (see Figure 1). Growth curves in fermentative (YPDextrose) and respiratory media (YPGlycerol) for yeast expressing each of the MRP truncations (in 2 replicates) are plotted besides WT controls not expressing any construct (grey lines); Curves are normalized by subtracting the minimal value of each to compensate for the background OD and then by plotting all curves only from the timepoint when the normalized OD reached 0.1 to compensate for different lag-times before the exponential phase; thus only the amplitudes and slopes of the growth curves can be compared but not the lag-times which are mostly attributed to inaccuracies of preparing the starting culture dilution.

**Fig. S6.**
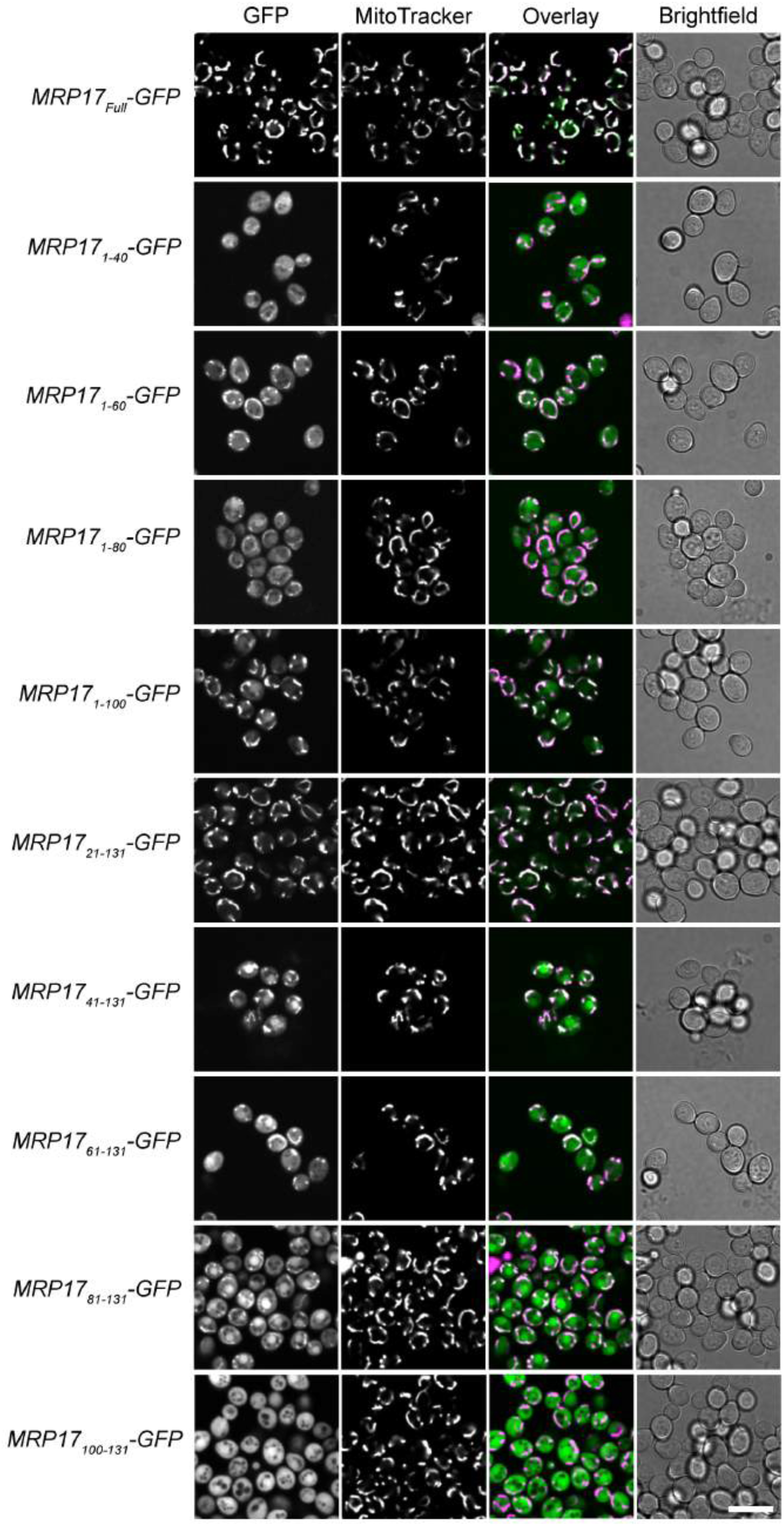
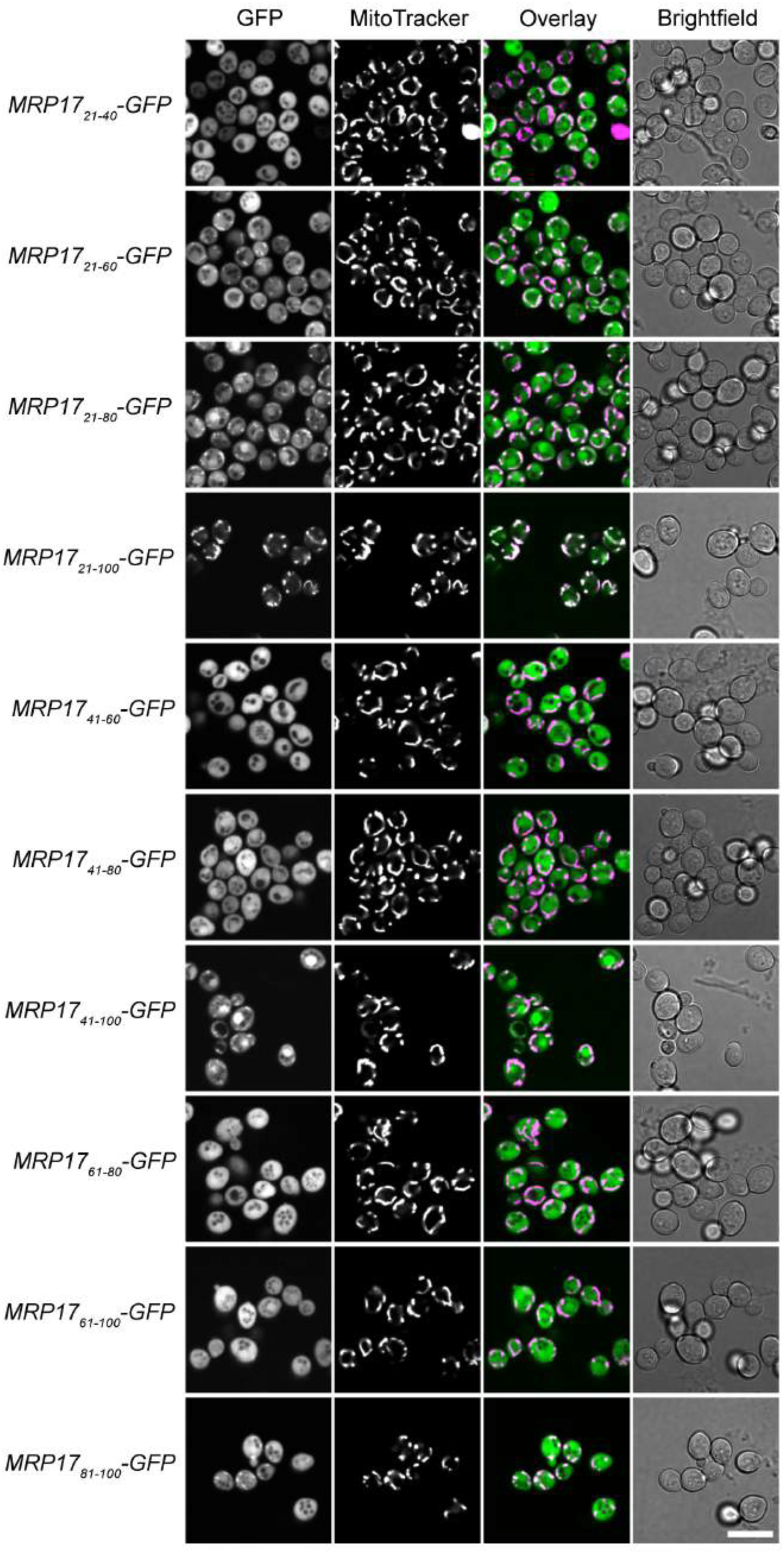
The ability of different Mrp17 truncations to target GFP to mitochondria. Fullsets of micrographs in different fluorescent channels for all the Mrp17 truncations including the ones shown in Fig. 2A. Scale bar 10 µm.

**Fig. S7.**
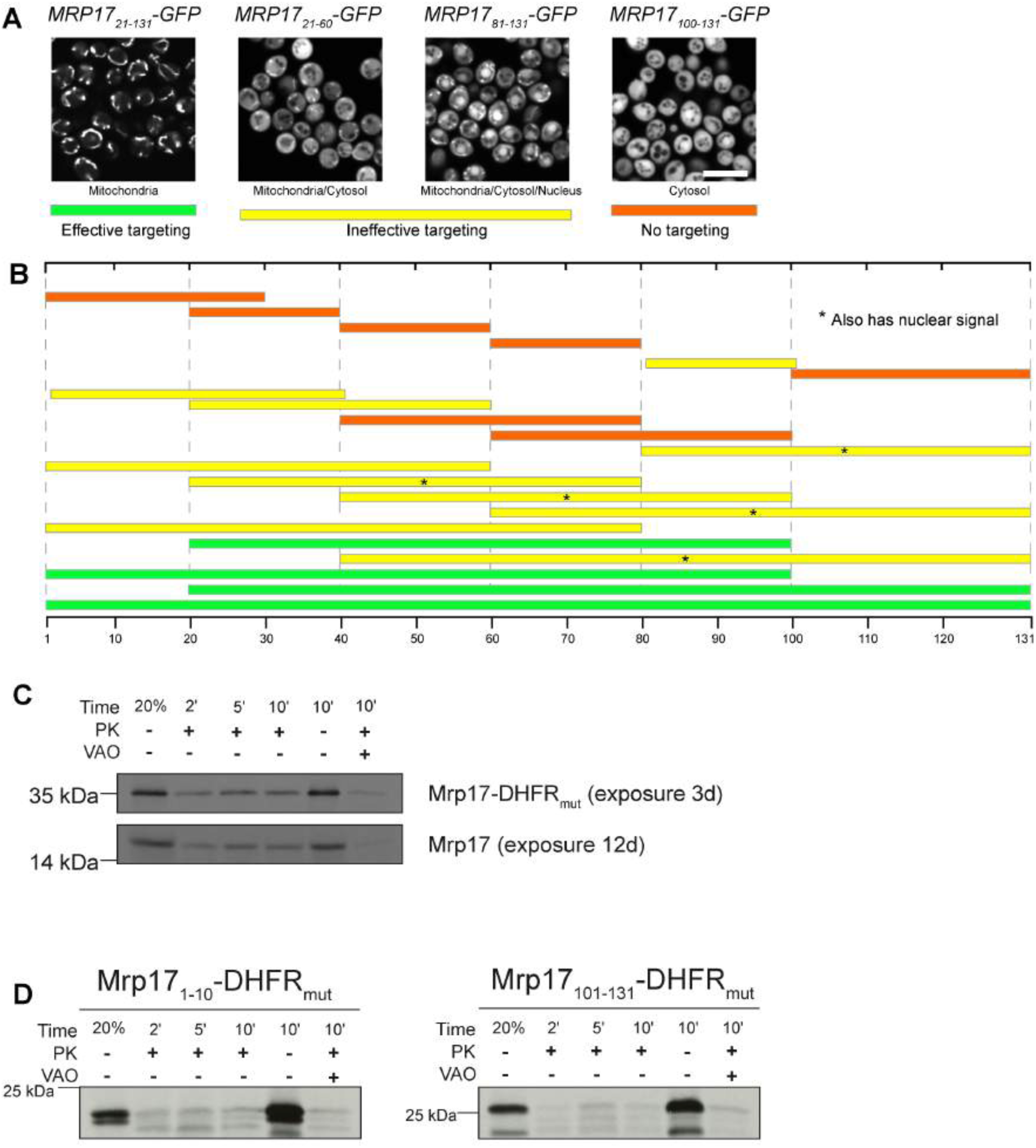
The non-canonical targeting and translocation signal of Mrp17 is located between amino acids 30 and 60. (Aand B) In vivo characterization of mitochondrial targeting capacity of different Mrp17 truncations fused to GFP: (A) GFP localization examples taken from Fig. S6 and demonstrating effective targeting with only mitochondrial GFP signal, ineffective targeting with mitochondrial GFP signal accompanied by strong cytosolic signal, ineffective targeting with additional nuclear signal, and no detectable mitochondrial targeting with exclusively cytosolic GFP. Scale bar is 10 µm. (B) localization summary of different Mrp17 truncations fused to GFP and colored according to the color-code for effective, ineffective, and no targeting introduced in panel (A), truncations additionally targeted to the nucleus are marked with asterisks. (C) Mrp17-DHFRmut is translocated into isolated mitochondria at the same rate as WT Mrp17 but gives better signal in the autoradiograph. (D) In vitro import assays for additional truncations of Mrp17 fused to DHFRmut not shown in Fig. 2B, import was performed as described in the legend for Fig.2.

**Fig. S8.**
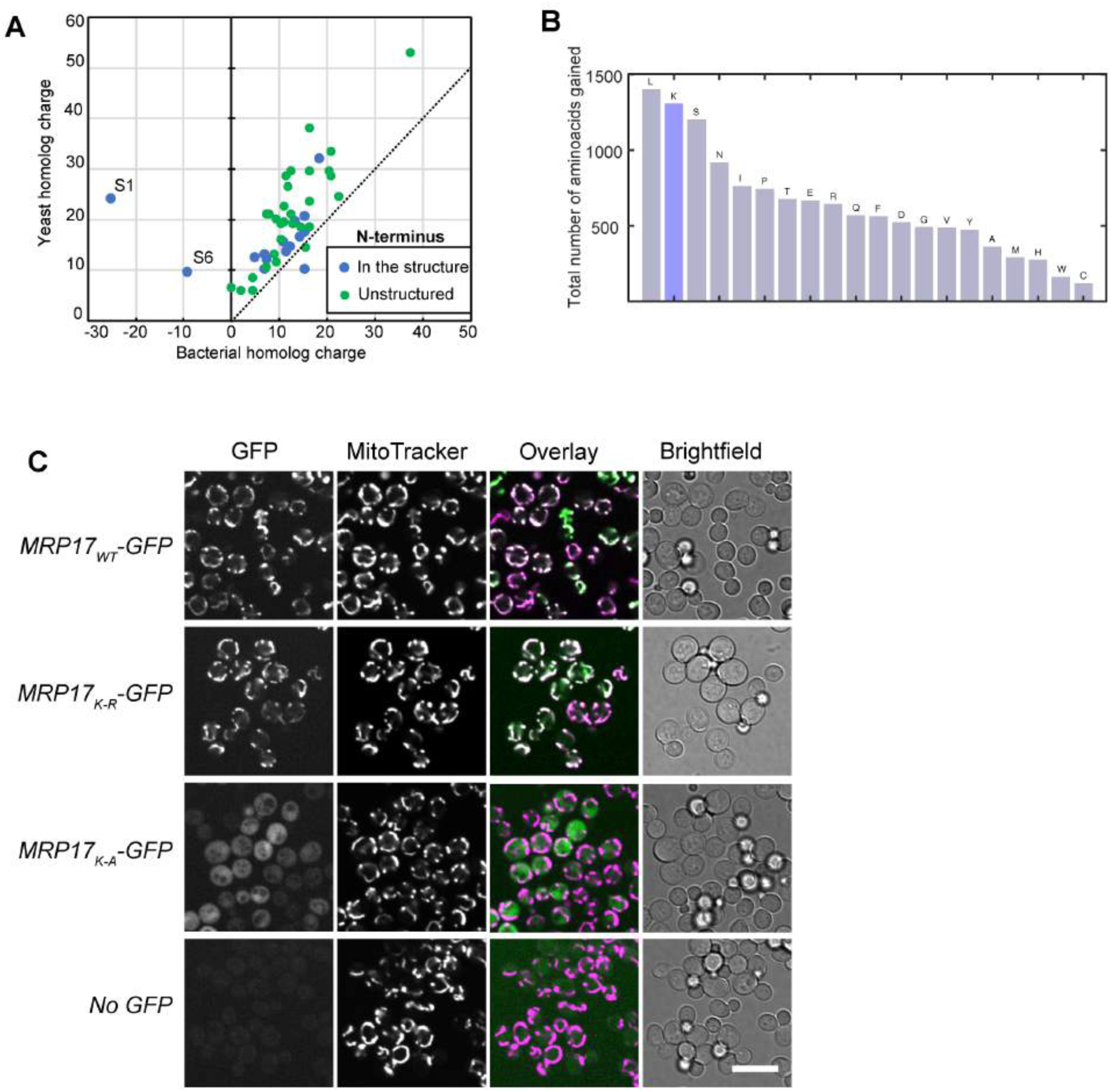
Mrp17 sequence features important for targeting and translocation to mitochondria. (A) Ribosomal proteins are positively charged and mitochondrial proteins acquired even more positive net charge compared to their bacterial homologs; (B) Total amino acid gain of MRPs (calculated as the difference between totalcount of each amino acid in all yeast MRPs, including mitochondria-specific, andall bacterial RPs) compared to bacterial RPs shows over-representationof lysines (K); (C) Lysines in Mrp17 are not important for mitochondrial targeting and can be substituted with arginines, same micrographs as in Fig. 3C shown in all channels beside micrographs of yeast not expressing any GFP (bottom row) as control for autofluorescence relative to cytosolic signal. All micrographs in the GFP channel are shown at the same contrast and brightness for comparison; Scale bar of all micrographs is 10 µm.

**Fig. S9.**
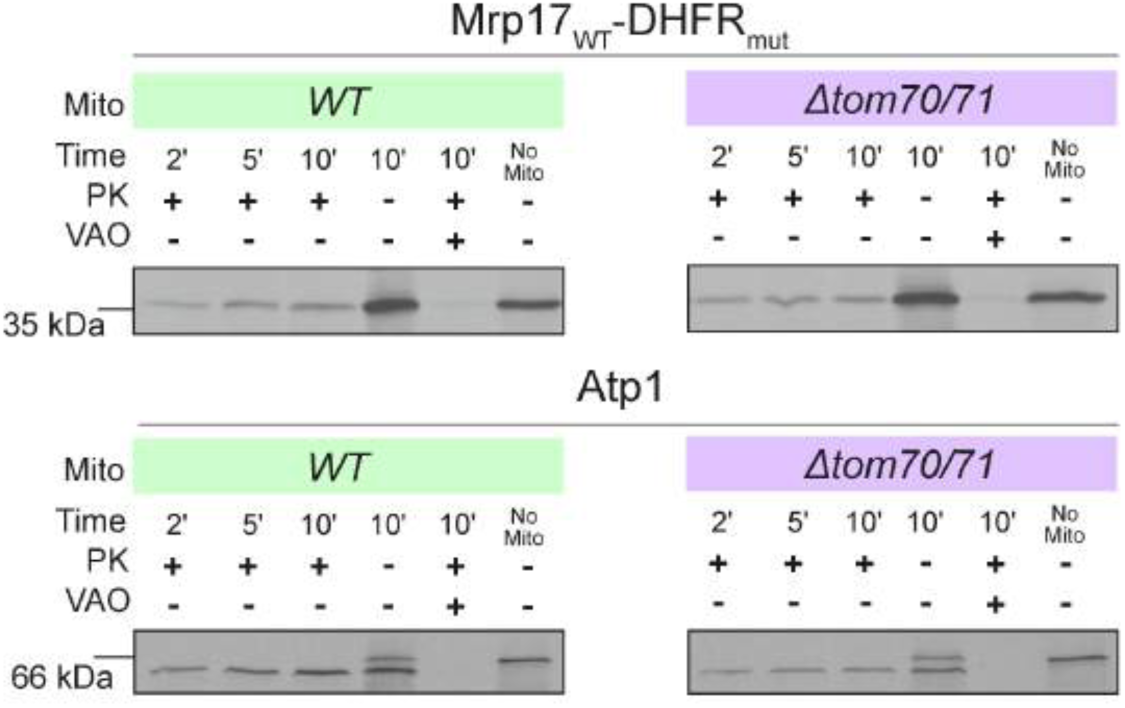
The translocation pathway of Mrp17 does not involve Tom70/71. *In vitro* translocation of Mrp17-DHFRmut and control protein Atp1 into mitochondria isolated from WT and a Δ*Tom70/71* mutant showing no Tom70/71-dependence for Mrp17 translocation. import was performed as described in the legend for Fig.2.

**Fig. S10.**
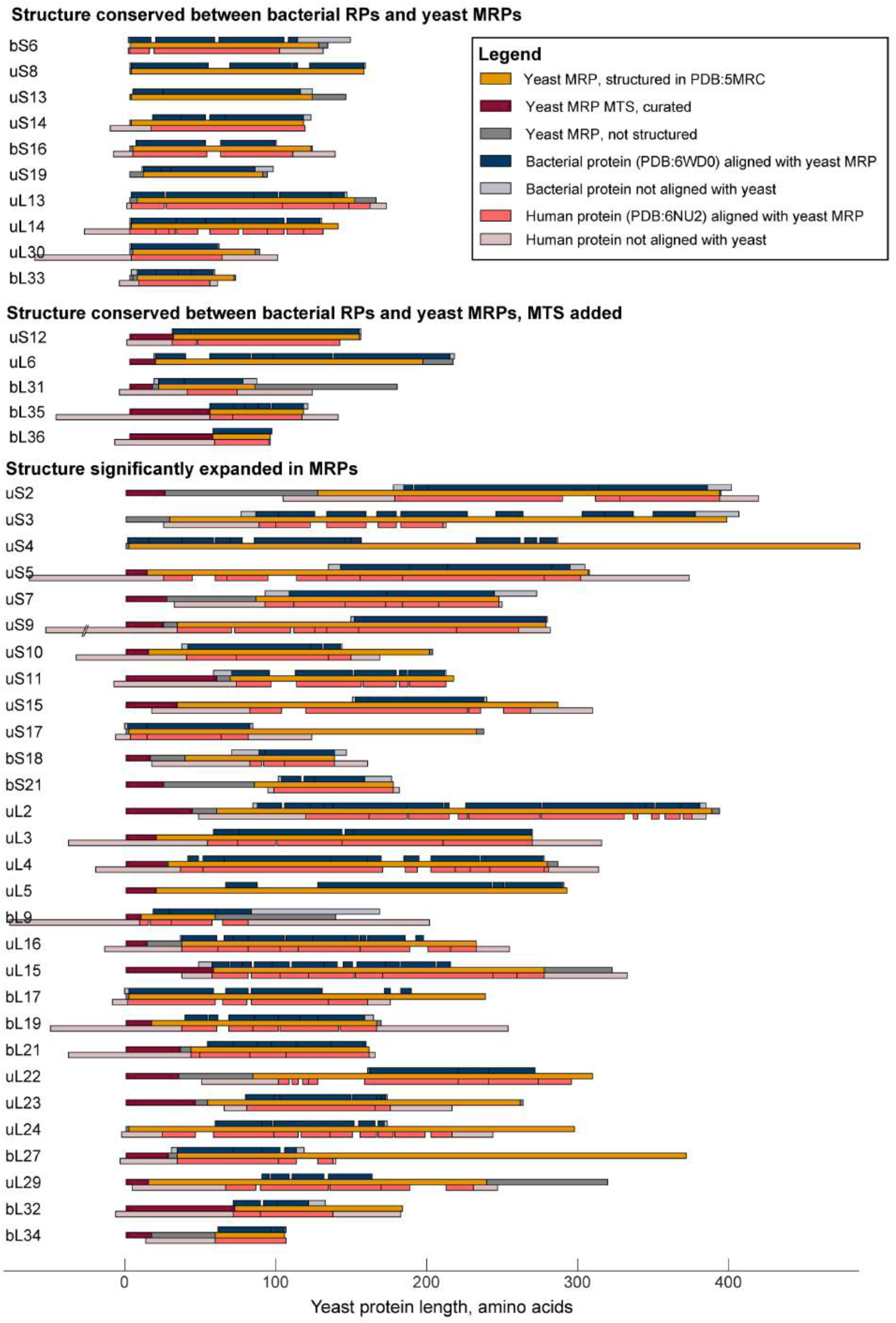
Structural alignments of yeast and human MRPs and their bacterial homologs. The structures of yeast MRPs that have bacterial homologs were aligned to their bacterial and human (when present) homologs using flexible structural alignment tool FATCAT (Li et al., 2020), and the resulting alignment was plotted relative to the length of yeast proteins, deletions in the yeast proteins relative to human and bacterial homologs are not plotted and the positions of corresponding insertions within human and bacterial sequences are indicated as solid black lines. MTS of yeast proteins were annotated according to Table S1. MPRs are grouped by structure conservation relative to bacterial homologs and then sorted by name.

**Fig. S11.**
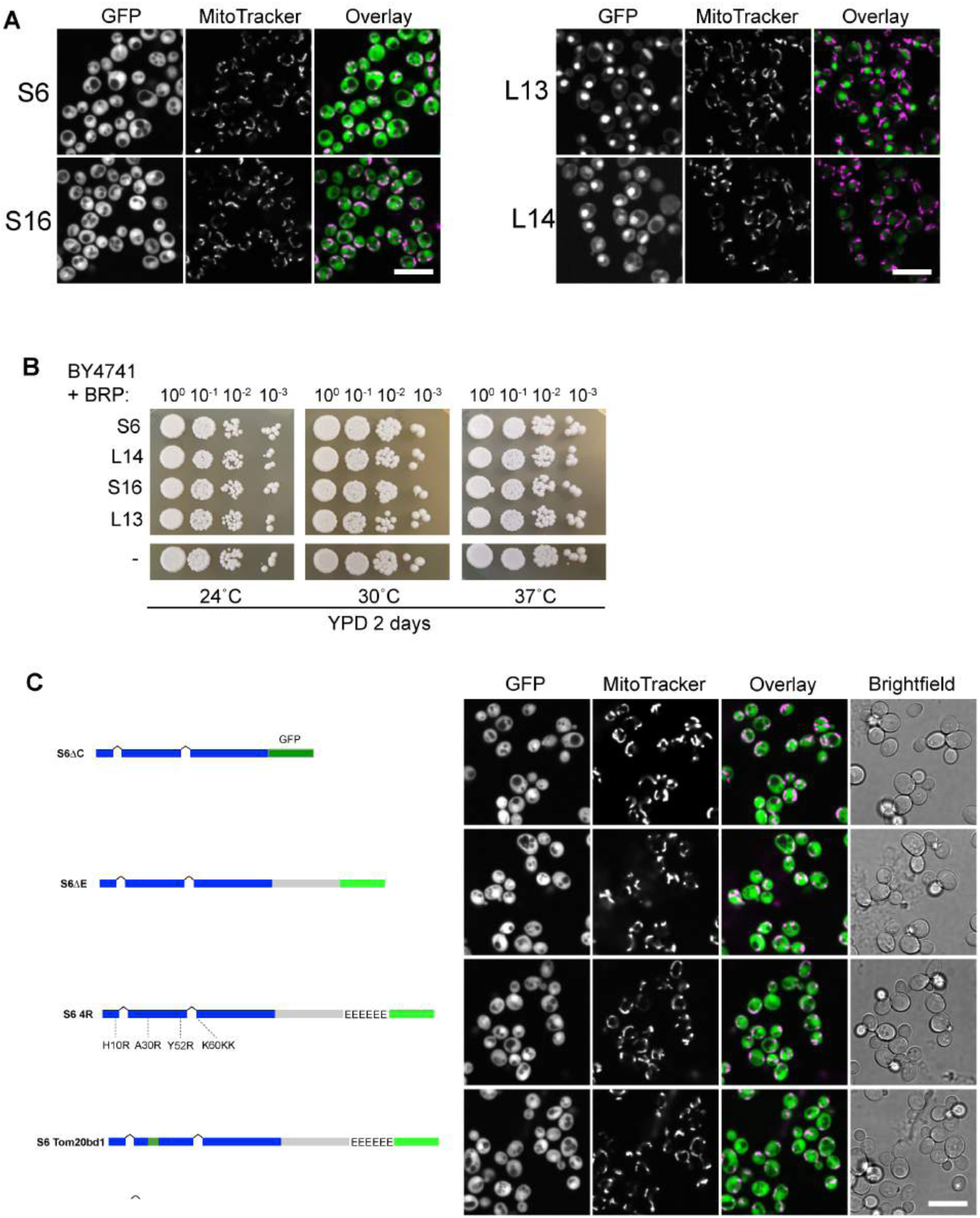
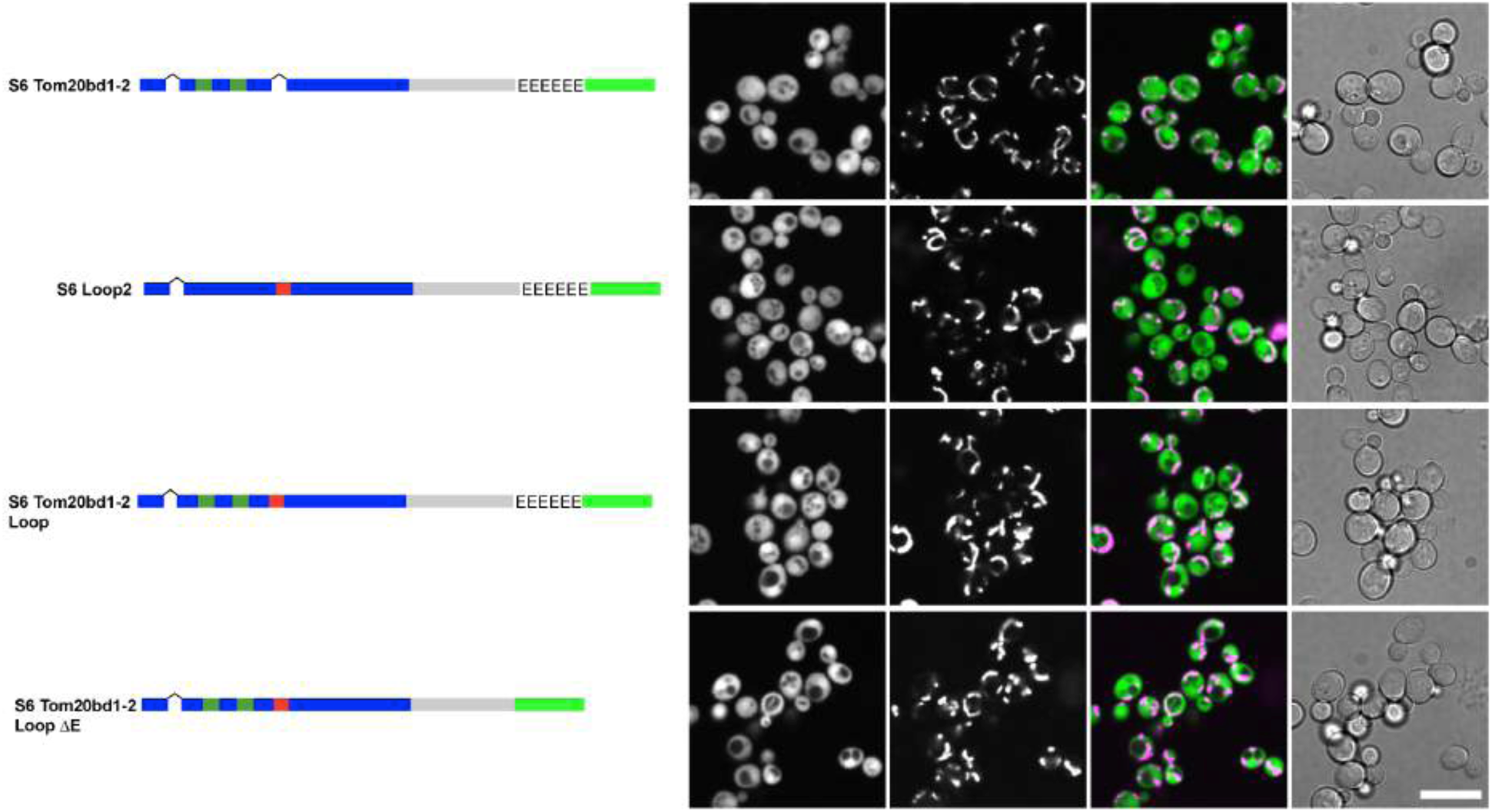
Comparison of Mrp17 and its bacterial homolog EcS6 in the structural context. (A) Expression of bacterial homologs of MRPs in yeast, same micrographs as in Fig. 5C shown in all channels. (B) Drop dilution growth assay for all the strains from panel A and WT control performed on rich fermentative media at different temperatures. (C) Schematic representations of chimeric constructs incorporating Mrp17 features into EcS6 (corresponds to schematic on Fig. 5D, top) and fluorescence micrographs of yeast expressing each respective construct. Scale bar in all panels is 10 µm.

**Fig. S12.**
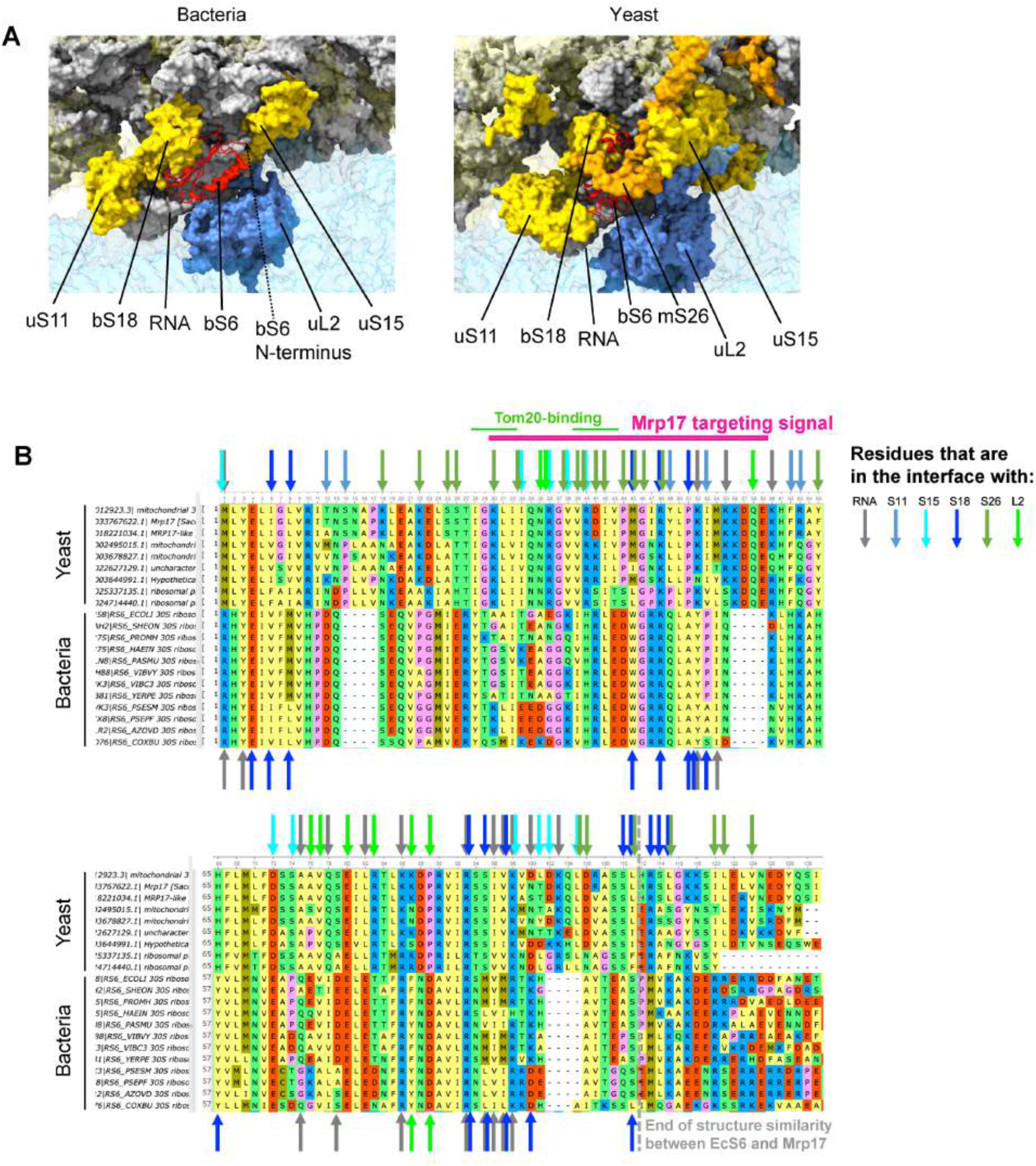
Mrp17 and EcS6 in the context of ribosome structures. (A) A comparison of the structural environment of the S6 protein in bacterial ribosome (PDB:6WD0, left) and yeast mitoribosome (PDB:5MRC, right). RNA is shown in grey, small subunit (SSU) proteins in yellow (highlighted are solid and others are transparent), S6 is red, all large subunit (LSU) components are in blue (highlighted are solid and others are transparent), mitochondria-specific protein mS26 is in orange. (B) Structural alignment of Mrp17 (first sequence) and its yeast homologs with EcS6 (first out of bacterial) and its bacterial homologs highlighting involvement of each amino acid in protein-protein and protein-RNA interfaces (arrows, see legend on the figure) in yeast mitoribosome(on top) and bacterial ribosome (in the bottom), amino acids are numbered relative to Mrp17, thus the N-terminal Met in bacterial proteins is not shown.

